# Repeated loss of function at HD mating-type genes and of recombination suppression without mating-type locus linkage in anther-smut fungi

**DOI:** 10.1101/2024.03.03.583181

**Authors:** Elise A. Lucotte, Paul Jay, Quentin Rougemont, Loreleï Boyer, Amandine Cornille, Alodie Snirc, Amandine Labat, Elizabeth Chahine, Marine Duhamel, Jacob Gendelman, Wen-Juan Ma, Roxanne Kaaren Hayes, Michael H. Perlin, Michael E. Hood, Ricardo C. Rodríguez de la Vega, Tatiana Giraud

## Abstract

A wide diversity of mating systems occur in nature, with frequent evolutionary transitions in mating-compatibility mechanisms. Basidiomycete fungi typically have two mating-type loci controlling mating compatibility, HD and PR, usually residing on different chromosomes. In *Microbotryum* anther-smut fungi, there have been repeated events of linkage between the two mating-type loci through chromosome fusions, leading to large non-recombining regions. By generating high-quality genome assemblies, we found that two sister *Microbotryum* species parasitizing *Dianthus* plants, *M. superbum* and *M. shykoffianum*, as well as the distantly related *M. scorzonarae*, have their HD and PR mating-type loci on different chromosomes, but with the PR mating-type chromosome fused with part of the ancestral HD chromosome. Furthermore, progressive extensions of recombination suppression have generated evolutionary strata. In all three species, rearrangements suggest the existence of a transient stage of HD-PR linkage by whole chromosome fusion, and, unexpectedly, the HD genes lost their function. In *M. superbum*, multiple natural diploid strains were homozygous, and the disrupted HD2 gene was hardly expressed. Mating tests confirmed that a single genetic factor controlled mating compatibility (i.e. PR) and that haploid strains with identical HD alleles could mate and produce infectious hyphae. The HD genes have therefore lost their function in the control of mating compatibility in these *Microbotryum* species. While the loss of function of PR genes in mating compatibility has been reported in a few basidiomycete fungi, these are the first documented cases for the loss of mating-type determination by HD genes in heterothallic fungi. The control of mating compatibility by a single genetic factor is beneficial under selfing and can thus be achieved repeatedly, through evolutionary convergence in distant lineages, involving different genomic or similar pathways.

## INTRODUCTION

Mating systems, corresponding to the degree of selfing/outcrossing, influence gene flow, genetic load and adaptability in natural populations^1, 2, 3, 4, 5, 6, 7^. A wide diversity of mating systems occur in nature, controlled by mating compatibility systems (e.g. separate sexes or mating types), with frequent evolutionary transitions between these systems^5, 8, 9, 10, 11^. In many animals and dioecious plants, different sexes are determined by sex chromosomes^12^. In hermaphrodite plants, mating compatibility is often controlled by a self-incompatibility locus, which prevents mating between identical alleles at this locus^13^. In fungi with mating types, gamete compatibility is controlled at the haploid stage, where only cells with different mating types can successfully mate^14,15^, mating types being determined by one or two loci^16^, the systems being then called uni- and bifactorial, respectively. There have been multiple transitions in fungi for changes in the number of loci controlling mating types, between one, two or even none, i.e., lack of discrimination for mating^8, 15^.

Recombination is typically suppressed at loci determining sexes and mating types, which ensures proper function in these complex phenotypes. Recombination suppression has furthermore often extended stepwise outward from these mating-compatibility loci in plants, animals and fungi^17,18, 19,20,21, 22, 23, 24^, and the reason why is still debated^23, 25^. It has long been accepted that recombination suppression extended on sex chromosomes because it was beneficial to link sexually antagonistic genes (i.e., with an allele being beneficial in only one sex) to the sex-determining locus^26^. However, little evidence could be found in favor of this hypothesis^27^, and it cannot explain the stepwise extension of recombination suppression on fungal mating-type chromosomes, as they lack sexual antagonism and other kinds of antagonistic selection^17, 23,28^.

Other hypotheses have therefore been developed, in order to account for the ubiquity of stepwise extension of recombination suppression on sex-related chromosomes: the neutral accumulation of sequence divergence proximal to the mating-compatibility locus that would reduce recombination rates^29^, the spread of transposable elements adjacent to the non-recombining mating-compatibility locus, together with their epigenetic marks^30^, or the selection of non-recombining fragments carrying fewer deleterious mutations than the population average while having their load sheltered by a permanently heterozygous allele^31-33^.

Following recombination suppression, Muller’s ratchet and Hill-Robertson effect^34,35^ lead to genomic degeneration, in terms of gene losses, weaker gene expression, lower frequencies of optimal codons, and transposable element accumulation^36,37, 38, 39,40, 41^. Rearrangements also rapidly accumulate^17,18,42^, so that recombination can be challenging to restore after the non-recombining fragments have become highly degenerated^31,33^.

Basidiomycete fungi typically have two mating-type loci controlling mating compatibility^16^: the PR locus, controlling pre-mating fusion with pheromone and pheromone receptor genes, and the HD locus, controlling post-mating dikaryotic growth with two homeodomain genes whose products heterodimerize to form an active transcription factor^43^. These two loci are usually on different chromosomes and may harbor multiple alleles. Some basidiomycetes in contrast are unifactorial, i.e. with a single segregating unit controlling mating type, which is caused most often by HD-PR linkage, as in the crop pathogen *Ustilago hordei*^44^ and human pathogens *Malassezia spp^45, 46^*, or more rarely by the loss of mating-compatibility role by the PR locus, as in the mushrooms *Coprinellus disseminatus*^47^ and *Volvariella volvacea^48^*. A loss of mating-compatibility role by the HD locus has only been observed so far in experimental mutants with their two HD genes fused^49^.

*Microbotryum* fungi (Basidiomycota) are plant-castrating pathogens, producing their spores in the anthers of diseased plants and therefore called anther-smut fungi. Most *Microbotryum* species specialized on one or a few host plants^50, 51, 52,53^. *Microbotryum* fungi are model organisms in as diverse fields as ecology, disease transmission, host specialization and reproductive isolation evolution, host-pathogen costructure, mating system and sex-related chromosome evolution^51, 53, 54, 55, 56, 57, 58, 59,60, 61, 62^. *Microbotryum* fungi mostly undergo selfing via intra-tetrad mating^58,63,64^, in which case carrying a single segregating mating-type locus is advantageous because it maximizes the odds of gamete compatibility^17^. *Microbotryum* fungi have in fact repeatedly evolved HD-PR linkage, with at least nine events of chromosomal rearrangements and recombination suppression between the two mating-type loci^17,18,39,41^. In addition, two non-sister *Microbotryum* species still have their mating-type loci on different chromosomes although each is linked to its centromere, resulting in the same odds of gamete compatibility as HD-PR linkage under intra-tetrad selfing^41,65^. Recombination suppression extended stepwise beyond mating-type loci in several lineages, forming evolutionary strata, i.e. fragments displaying decreasing differentiation with increasing distance to mating-type loci along the ancestral gene order^17,18,41^.

The non-recombining regions on both mating-type chromosomes in *Microbotryum* fungi display signs of genomic degeneration, as neither ever recombines. For example, polymorphism is common for haploid sporidia of one or the two mating types failing to grow *in vitro* in some *Microbotryum* species, likely due to deleterious mutations linked to mating-type loci^58,66-68^. Indeed, with time since recombination suppression, these mating-type chromosomes accumulate transposable elements, non-synonymous mutations, gene losses, rearrangements and non-optimal codons ^18,39,40,42,69^. Transposable elements accumulate rapidly following recombination suppression, and some specific transposable element families expanded preferentially on *Microbotryum* mating-type chromosomes, such as *Helitrons* and *Copia/Ty3* elements^69,70^. These patterns are especially pronounced in species with linked HD and PR loci and extensive suppression of recombination, in contrast to the genomes retaining the ancestral state of unlinked mating types, that exhibit lesser severity of degenerative processes^18,39,40,42,69.^

Surveys of the diversity within *Microbotryum* fungi continue to yield novel insights, and preliminary analyses suggested that the species parasitizing *Dianthus* plant species carried unlinked HD and PR loci, yet displaying very high transposable element content compared to other *Microbotryum* species^70^. Using two *Microbotryum* species on *Dianthus*, our aim was therefore to characterize the mating-type chromosomes of these species, elucidate whether the mating-type loci display localized recombination suppression, and if so, whether the mating-type loci were linked together or to their respective centromeres. We found in both species a large non-recombining mating-type chromosome issued from the fusion of the entire ancestral PR chromosome and a part of the ancestral HD chromosome but, surprisingly, not HD locus itself. Genome comparison indicated that the translocation occurred via a transient stage of whole chromosome fusion, a few rearrangements and then separation of a part of the HD chromosome arm. Remarkably, homozygosity was observed at the HD genes in natural populations, indicating that they were not involved anymore in mating-type determination, and one of the HD2 alleles appeared disrupted in *M. superbum*. This novel observation among basidiomycete fungi was also found in a distant congeneric species, *M. scorzonerae*, with unlinked PR and HD loci, but homozygosity and non-functionalization of HD genes, and rearrangements between ancestral HD and PR chromosomes. These results demonstrate the potential for genetic innovation in compatibility systems that directly influence how mating system variation is achieved in nature. In addition, we found multiple evolutionary strata in the three species, i.e., young extensions of recombination suppression on the mating-type chromosomes.

## RESULTS

### A large non-recombining PR chromosome including an arm of the ancestral HD chromosome but not the HD genes

We obtained high-quality genome assemblies from one strain from each of two *Microbotryum* species parasitizing *Dianthus* plants: *M. superbum* (aka MvDsp2^52^, strain 1065) and *M. shykoffianum* (aka MvDsp1^52^, strain c212). The phylogenetic tree based on 2,391 single-copy orthologs representing 5.2 mb of nucleotide sequences placed them as sister species as expected (Figure 1). In all four haploid genomes, the PR and HD genes were found on different contigs (Figures 2 and S1-3) and with telomeric repeats identified on almost all of these mating-type contig edges (Figure S4-6), showing that the two mating-type loci are located on different chromosomes. We compared these mating-type chromosomes to those of *M. lagerheimii*, inferred to be a good proxy for the ancestral gene order of *Microbotryum* mating-type chromosomes^17,41^. We found the PR gene to be located in both *M. superbum* and *M. shykoffianum* on a chromosome corresponding to the fusion of the ancestral PR chromosome and a large part of the ancestral HD chromosome. The remaining part of the ancestral HD chromosome, carrying the HD gene, was found on a separate chromosome (Figures 2 and S1-4). This new PR chromosome was non-recombining across most of its length, as it was widely rearranged between species and between a_1_ and a_2_ chromosomes in both *M. superbum* and *M. shykoffianum* (Figures 2 and S1-5). Only small regions were collinear at both edges of the PR chromosome, corresponding to recombining pseudo-autosomal regions (PAR, Figures S4-S5). The centromere was identified within the non-recombining region and telomeres at the edges of the new PR and HD chromosomes (Figures S4-S5). The separate chromosome carrying the HD genes was very small in both species, and collinear between the two mating types as well as between species (Figures 3 and S1-5), suggesting ongoing recombination.

**Figure 1:**
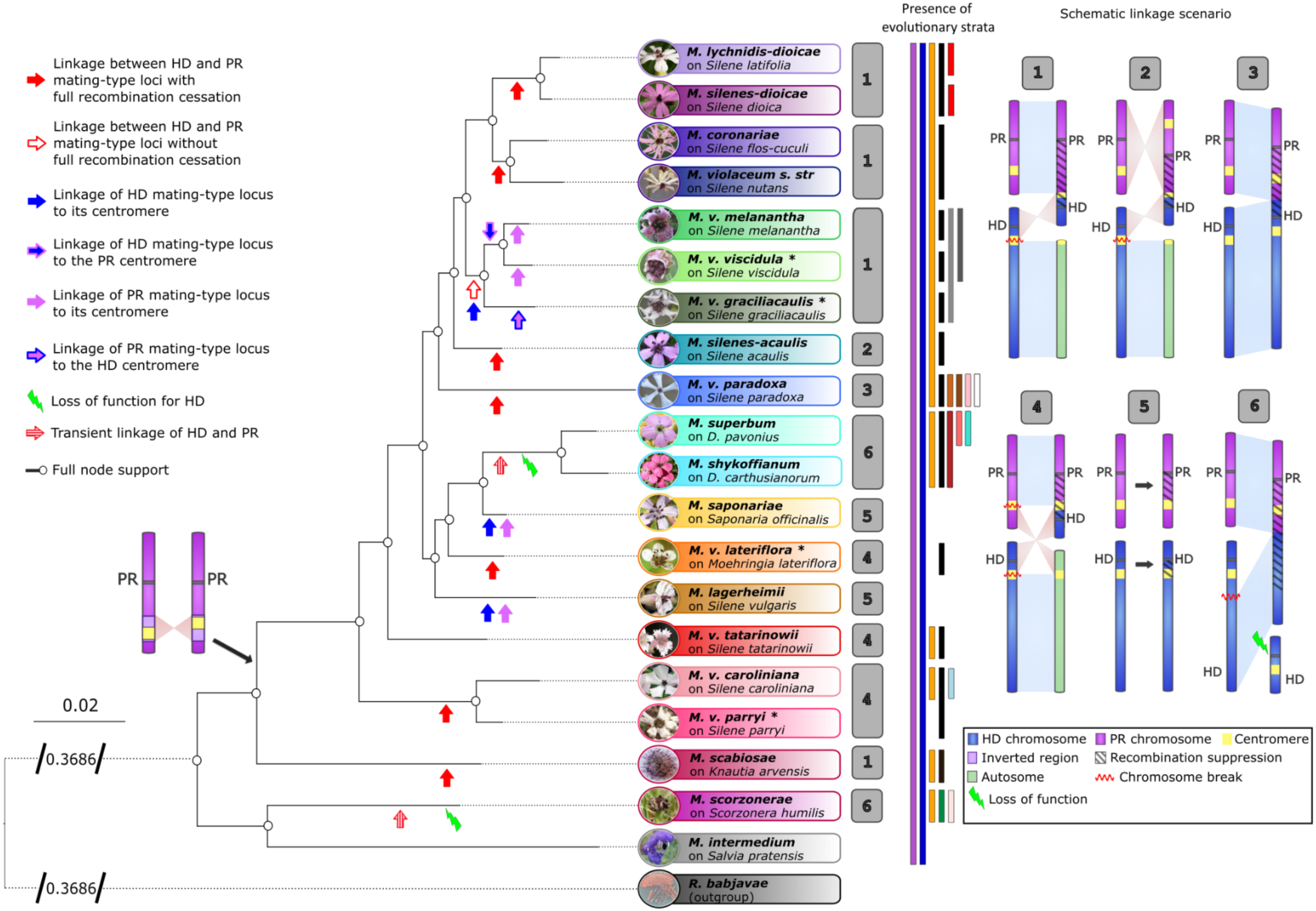
Species tree with pictures of diseased plants parasitized by the different *Microbotryum* species and showing chromosomal arrangements of the mating-type chromosomes compared to the ancestral state. Arrows of different colors on the tree branches indicate mating-type locus linkage, or their linkage to centromeres, or loss of function. Colored bars at the right of the phylogeny indicate the presence of evolutionary strata suppression recombination beyond mating-type loci. Pictures are from Michael E. Hood and Julian Woodman.

**Figure 2:**
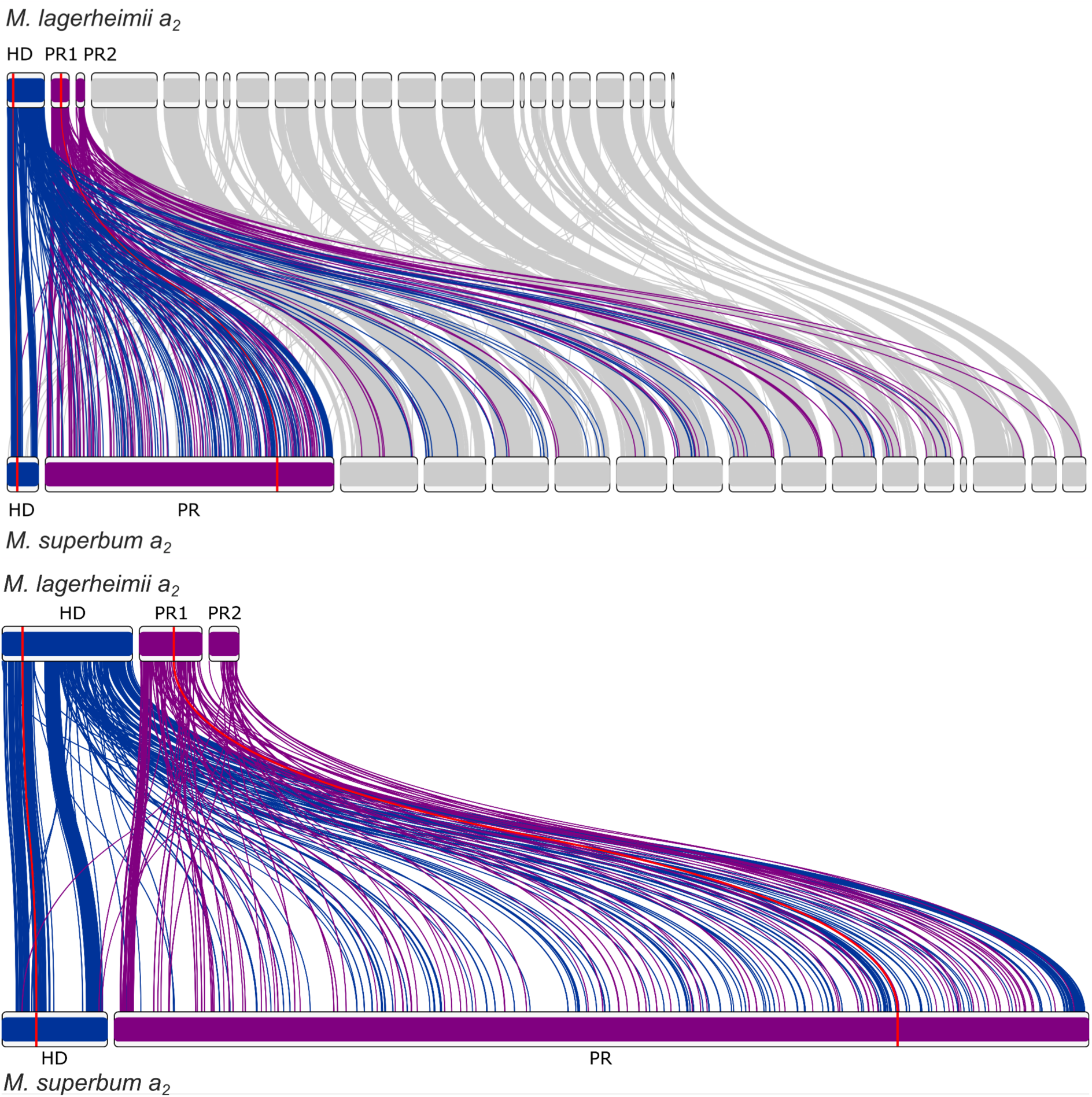
Synteny and rearrangements between the a_1_ genomes of *Microbotryum superbum* and *M. lagerheimii*, with all contigs (top) or only the mating-type chromosomes (bottom). The *M. lagerheimii* genome represents a proxy for the ancestral state before recombination suppression and with two separate mating-type chromosomes. The HD mating-type chromosome from *M. lagerheimii* is represented in blue and the PR mating-type chromosome in purple (splitted into two scaffolds), with links to orthologous genes to the *M. superbum* mating-type chromosomes. The positions of the HD and PR mating-type genes are indicated in red and by red links between the two genomes. The PR chromosome shows a substantial increase in size and a chaos of rearrangements.

**Figure 3:**
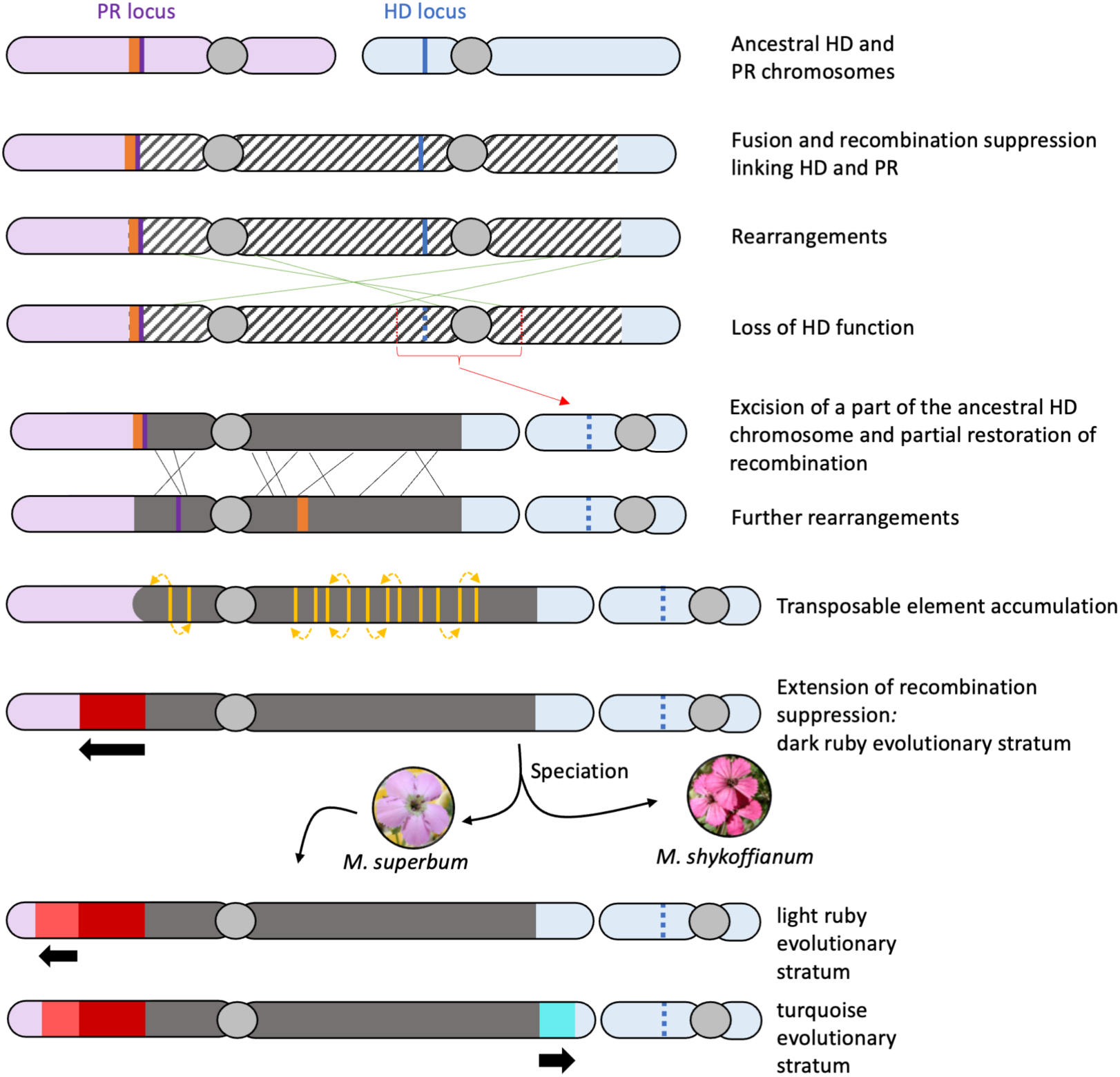
Inferred scenario for the evolution of the mating-type chromosomes in *Microbotryum superbum* and *M. shykoffianum.* Pictures of the diseased plants parasitized by the two species are shown. Recombination suppression is figured by colors corresponding to the different evolutionary strata.

In both *M. superbum* and *M. shykoffianum*, a few genes present on the HD chromosome part carrying the HD genes in *M. lagerheimii* were however found in the non-recombining region of the PR chromosome, and conversely (genes colored in green in Figures 3-4 and S4). This indicates an ancient fusion between the whole PR and HD chromosomes, followed by a few genomic rearrangements, before an excision of a part of the ancestral short HD chromosome arm (Figure 3). Such an excision may have occurred in a single step or by chromosome fissions followed by a new fusion. Some of the genes that were found reshuffled between the ancestral PR and HD chromosomes were different between *M. superbum* and *M. shykoffianum*, which may suggest that the fission of the current small HD chromosome occurred independently in the two species, after their speciation.

**Figure 4:**
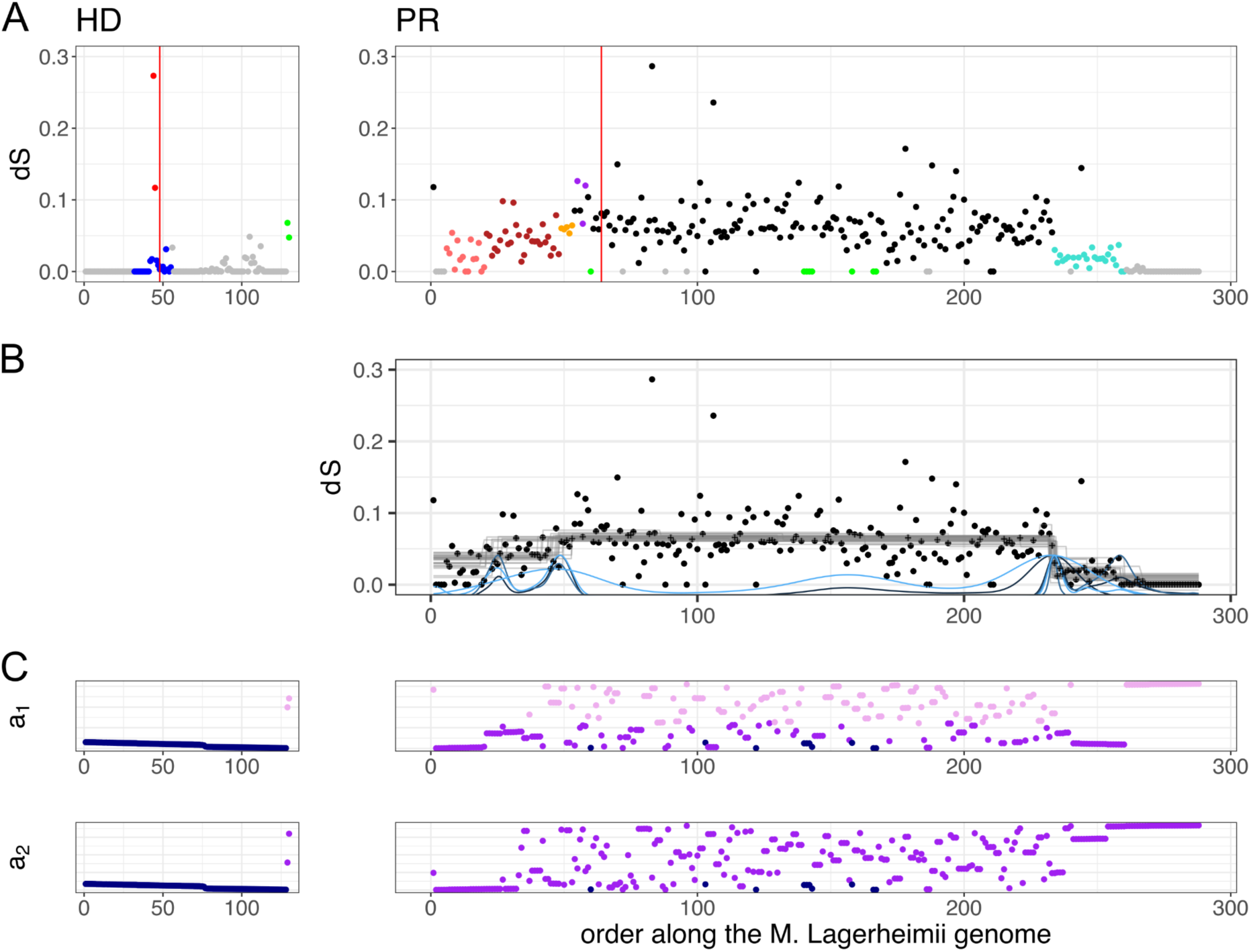
Differentiation and rearrangements between the mating-type chromosomes in *Microbotryum superbum*. **A)** Per-gene synonymous divergence (d_S_) plotted along the ancestral gene order (taking as proxy the gene order along the *M. lagerheimii* mating-type chromosomes), the HD chromosome at left and the PR chromosome at right; the points are colored according to their evolutionary stratum assignment: turquoise, light and dark ruby for the *M. superbum* specific evolutionary strata, black for the evolutionary stratum shared by *M. superbum* and *M. shykoffianum,* blue, orange and purple for the ancient evolutionary strata shared by most *Microbotryum* species. The pseudo-autosomal regions (PARs) are in grey. The green points correspond to genes that were ancestrally on the PR chromosome but found in the HD chromosomes in *M. superbum* or reciprocally. **B)** Change-point analysis identifying changes in mean dS levels. **C)** Rearrangements compared to the ancestral gene order figured by plotting the gene rank in the current gene order (b1 in the a_1_ genome and b_2_ in the a_2_ genome) as a function of the gene rank in the ancestral gene order.

Nucleotide polymorphism data further indicated that the PR chromosome was mostly non-recombining and that the HD genes were not linked to the PR gene in the *Microbotrum* fungi parasitizing *Dianthus* plants. Using SNPs on 149 Illumina genomes collected on various *Dianthus* plants in western Europe, high levels of LD were found across most of the PR chromosome, while LD levels were much lower between HD and PR contigs (Figure S7). The LD values within the HD chromosome, as well as against the PR chromosome, were a bit higher than those within autosomes, but were similar as those between the HD chromosomes and autosomes, thus not indicating linkage.

The high per-gene synonymous divergence (d_S_) between alleles on the a_1_ and a_2_ mating-type chromosomes (Figures 4-5) further confirmed the lack of recombination along most of the PR chromosome, while the HD chromosome appeared recombining. Indeed, because selfing in *Microbotryum* anther-smut fungi yields high homozygosity levels, recombining region shows zero d_S_ values between alleles in recombining regions such as autosomes, pseudo-autosomal regions of the PR chromosome and in the HD chromosome (Figures 4-5). In contrast, the rearranged region of the PR chromosome displayed high d_S_ values in both *M. superbum* and *M. shykoffianum* (Figures 4-5). Following previous studies on *Microbotryum* fungi, the region initially linking the PR and HD genes was called the black stratum (Figures 4-5), although this stratum does not link the PR and HD mating-type loci any more due to the proposed fission of the HD-containing chromosome arm (Figure 3). This black stratum evolved before the divergence of the *M. superbum* and *M. shykoffianum* lineages, as 73% of the genes showed full trans-specific polymorphism between these two lineages (Figure 6): the alleles clustered according to a_1_ or a_2_ mating type rather than species in gene genealogies, indicating recombination suppression before speciation. We found footprints of previously inferred evolutionary strata (i.e. the blue, purple and orange strata; Figures 4-5), that were shown to have evolved around the HD and PR genes ancestrally in the *Microbotryum* clade^17,18,23^.

**Figure 5:**
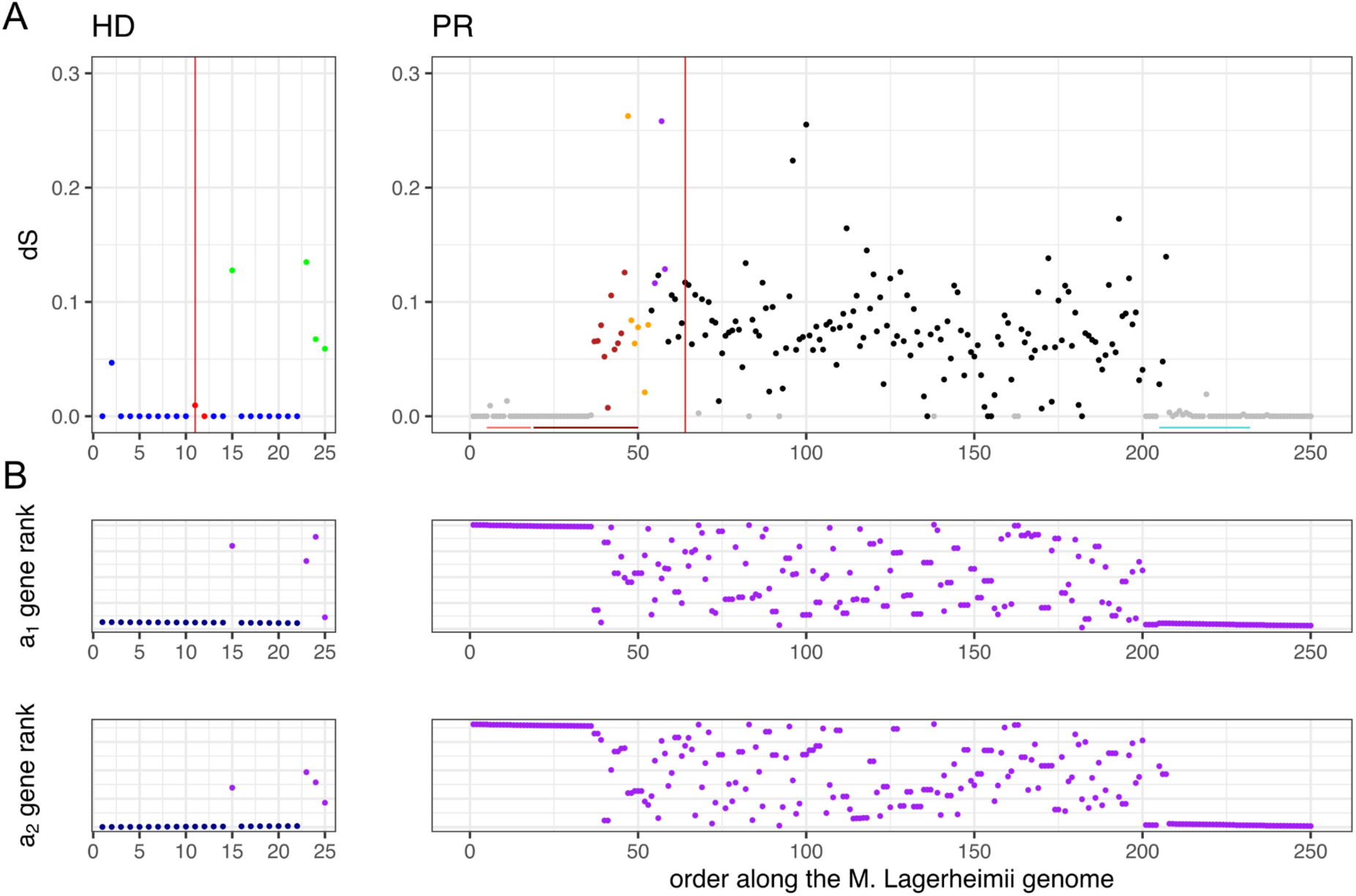
Differentiation and rearrangements between the mating-type chromosomes in *Microbotryum shykoffianum*. **A)** Per-gene synonymous divergence (d_S_) plotted along the ancestral gene order (taking as proxy the gene order along the *M. lagerheimii* mating-type chromosomes), the HD chromosome at left and the PR chromosome at right. The points are colored according to their evolutionary stratum assignment: turquoise, light ruby and half of the dark ruby region for the *M. superbum* specific evolutionary strata, black and the other half of the dark ruby for the evolutionary stratum shared by *M. superbum* and *M. shykoffianum,* blue, orange and purple for the ancient evolutionary strata shared by most *Microbotryum* species. The pseudo-autosomal regions (PARs) are in grey. The position of the *M. superbum* specific evolutionary strata are indicated by color bars at the bottom. The green points correspond to genes that were ancestrally on the PR chromosome but found in the HD chromosomes in *M. superbum* or reciprocally. **B)** Rearrangements compared to the ancestral gene order figured by plotting the gene rank in the current gene order (b_1_ in the a_1_ genome and b_2_ in the a_2_ genome) as a function of the gene rank in the ancestral gene order.

**Figure 6:**
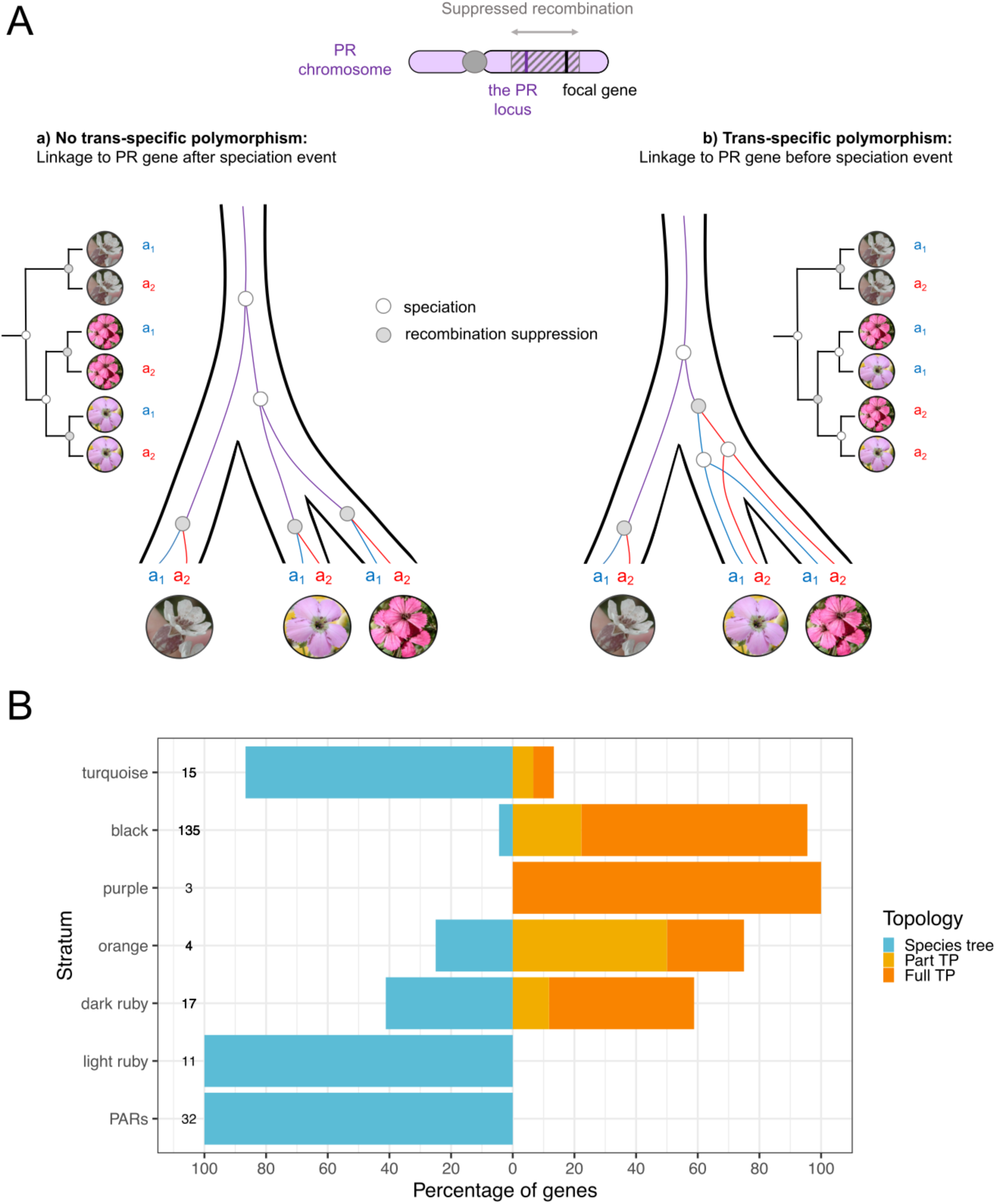
Trans-specific polymorphism (TP) of genes present along the mating-type chromosomes in *Microbotryum superbum*, considering also *M. shykoffianum* and *M. lagerheimii.* A) Schematic representation of trans-specific polymorphism in genes completely linked to mating-type loci by suppressed recombination, allowing to assess whether recombination cessation occurred before or after speciation, here between *M. lagerheimii, M. superbum* and *M. shykoffianum*. **B)** Percentage of genes within each evolutionary stratum with the following genealogy pattern: full trans-specific polymorphism (i.e., across both *M. superbum* and *M. shykoffianum*), partial trans-specific polymorphism (i.e., with only one allele showing clustering by mating type and the other by species, or with one allele missing, i.e. not informative), and no trans-specific polymorphism or unresolved. The total number of genes analyzed per stratum is indicated at the left of each barplot. The PARs (pseudo-autosomal regions) correspond to the PR chromosome PARs pooled together.

In the *M. shykoffianum* genome, the HD chromosome in the a_2_ assembly was much shorter than in the a_1_ assembly, while still carrying the HD gene (Figure S4-5). The mapping of diploid reads on the a_1_ plus a_2_ genome assemblies showed double coverage in the missing regions of the HD chromosome, indicating the collapse of the two homologous chromosomes in the corresponding regions in the a_1_ assembly (Figure S8). The coverage was however not doubled in a small region at the edge of one pseudo-autosomal region, and the corresponding genomic region was not found in the a_2_ genome assembly, suggesting that this small region may be genuinely missing in the a_2_ genome (Figure S8).

### Loss-of-function in mating-type determination of HD genes and lack of PR-HD linkage established by segregation analyses

The alleles of the two tightly-linked HD genes (HD1 and HD2) appeared functional in the two haploid genomes of *M. shykoffianum* and in the a_1_ genome of *M. superbum*, but not in the a_2_ genome of *M. superbum*. The HD2 gene indeed appeared separated into two different protein coding sequences (CDS) in the a_2_ genome in *M. superbum,* with an early stop codon and a new start codon (Figure 7). One of the two novel CDS appeared expressed during mating conditions, but not *in planta*, and we detected no expression of the HD2 gene in mating conditions or when cultured as haploid. This may be because the strain is homozygous or because the a1 allele of HD2 is not expressed. In contrast, in *M. lychnidis-dioicae* (with non-recombining mating-type chromosomes) and *M. intermedium* (with almost fully recombining mating-type chromosomes) both HD genes are upregulated under mating or *in planta* conditions (Figure 7). Analyses of non-synonymous substitution proportions based on gene genealogies with all available species (Figure 1) indicated significant positive selection in the two new CDS, suggesting neofunctionalization (significant test for selection intensification: K = 22.64; p = 0.001, LR = 11.37).

**Figure 7:**
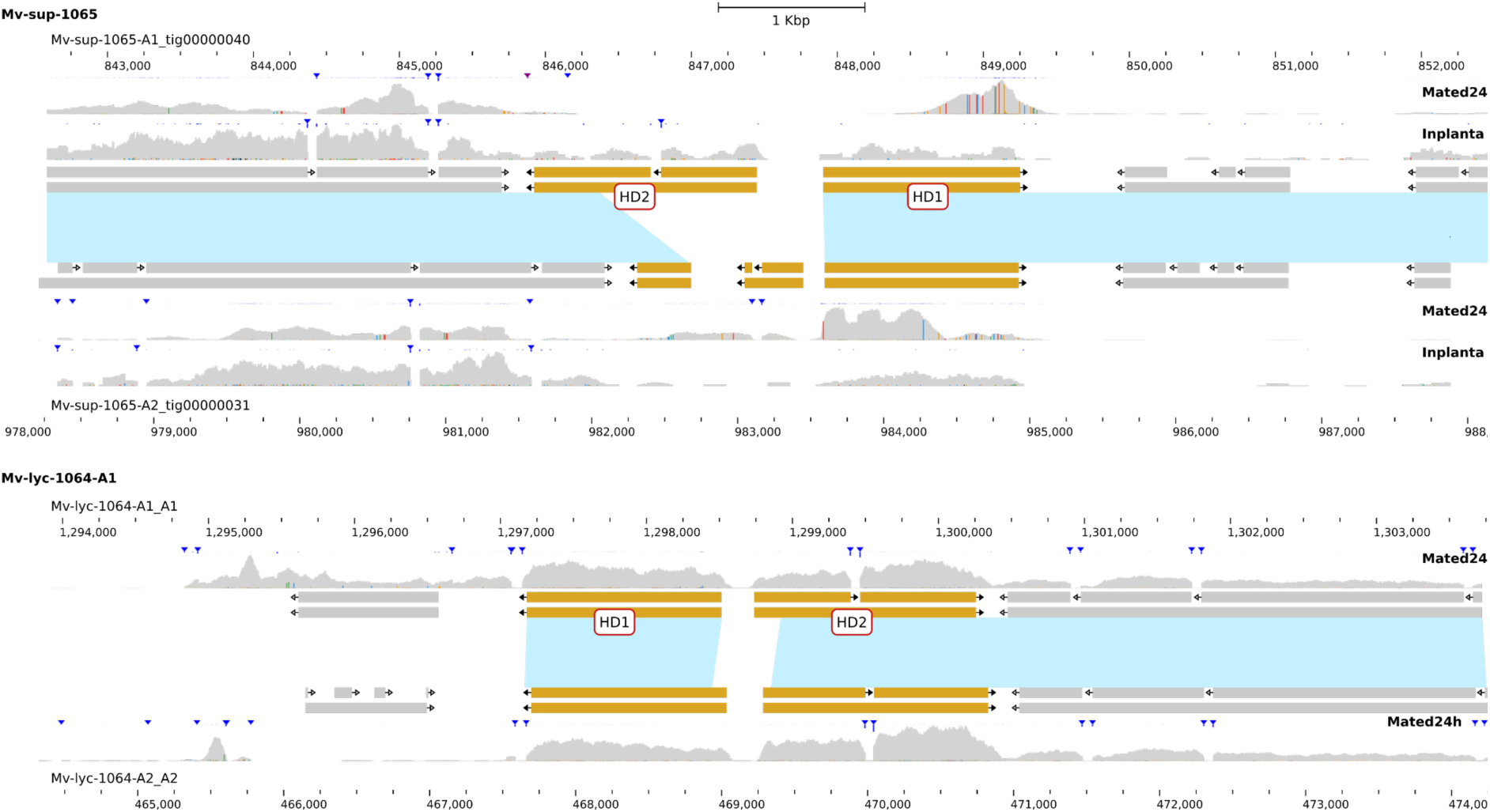
Structure and expression of the HD genes in *Microbotryum superbum* (A) and *M. lychnidis-dioicae* (B). In the middle of each panel, the coding sequences of the genes present in the a_1_ HD chromosome and the a_2_ HD chromosome are represented by rectangles (upper rectangles correspond to the coding sequence features and lower rectangles to the transcript features), and the direction of the transcription is indicated by an arrow. The HD2 and HD1 genes are represented in yellow boxes. The blue areas represent the syntenic regions between the a_1_ and a_2_ sequences. Above and below the rectangles, the level of expression is represented and annotated according to the condition of the RNAseq extraction (mated 24h or *in planta*).

Mating tests confirmed that HD genes have lost their mating-type determination function in *M. superbum*. We isolated the four meiotic products of 12 full tetrads from a strain collected on *D. pavonius* in the same site as the reference strain. *Microbotryum* produces linear tetrads allowing assessment of first versus second division segregation^58,71^. We tested whether each of these haploid cell lineages would mate with tester haploid strains from one of the tetrads that carried a_1_ or a_2_ PR alleles, respectively, as determined by PCR^72^. The typical situation in heterothallic basidiomycete fungi is that i) only haploid cells carrying different PR alleles can fuse and ii) only haploid cells carrying different HD alleles can produce hyphae when mated. In all 12 tetrads but one, two of the four meiotic products conjugated with the a_1_ tester and the other half with the a_2_ testers, as expected, with PR alleles segregating at the first meiotic division, consistent with linkage of the PR gene to its centromere shown above. Supporting the view that the HD genes no longer have a role in mating-type determination, all meiotic products compatible for conjugation with a given mating tester also produced hyphae. If HD genes were still involved in mating-type determination, only half of conjugated pairing would produce hyphae due to independent segregation/assortment of PR and HD mating type loci inferred from genomic assemblies.

Additionally, PCR tests confirmed that the 12 meiotic tetrads analysed in *M. superbum* resulted from a field-collected diploid parental genotype that contained identical HD alleles, which indicates the lack of mating-type role of HD genes. We indeed designed primers that would specifically amplify only the allele present in the a_1_ or a_2_ genome of the *M. superbum* 1065 strain (Table S1). The primers for the HD1 and HD2 genes gave PCR products only with the primer pair designed to amplify the alleles in the a_2_ genome of the *M. superbum* 1065 strain, and in all four meiotic products of all tetrads, indicating homozygosity of the diploid parent. Sanger sequencing of the amplicons confirmed that all meiotic products were identical at the HD genes. Therefore, heterozygosity at the HD genes was not required any more in these strains for successful mating and hyphae production. We also inoculated *D. chinesis* in the greenhouse and showed these homozygous pairs could cause disease (5 out of 24 inoculated plants, as typical under inoculations). In one tetrad, two meiotic products produced hyphae alone without any tester strains and the PCR indicated that these sporidia contained the two PR alleles, likely following abnormal meiotic segregation (i.e. non-disjunction), as previously reported in *Microbotryum* fungi (back then called *Ustilago violacea*)^73^. This sporidial line however did not produce disease upon plant inoculation.

We further pulled out two tetrads from each of six additional field-collected *M. superbum* strains. Two strains were homozygous for the HD genes, confirming that mating can occur between cells with identical HD alleles; the other strains were heterozygous. Across tetrads, the PR alleles were not always associated to the same HD alleles, which confirms that the HD and PR loci are unlinked in this species (Table S2).

The HD1 and HD2 coding sequences in *M. shykoffianum* c212 were not disrupted but were 100% identical between the a_1_ and a_2_ genomes, while the PR alleles showed high differentiation and trans-specific polymorphism as expected. This reinforces the view that the HD genes can be homozygous in *Microbotryum* strains from *Dianthus* species, and that, therefore, HD genes are not involved anymore in mating-type determination.

Polymorphism analysis of the HD genes based on Illumina sequencing confirmed that they could be homozygous in natural strains. We used SNPs called in the 149 Illumina genomes against the a_1_ *M. superbum* reference plus the HD and PR chromosomes from the a_2_ genome. The genomes were obtained by spreading diploid teliospores on culture medium, where they undergo meiosis and haploid meiosis product multiply. In 23 strains, we found only significant mapping on the a_1_ PR chromosome and in 14 strains only on the a_2_ PR chromosome. This is likely due to haplo-lethal alleles on the PR chromosome, with gene disruptions that are lethal at the haploid stage due to degeneration following recombination suppression, preventing growth of haploid sporidia on our culture plates, as identified in several *Microbotryum* species^58,66,68^. Among the 62 strains for which we were confident that they carried the two PR chromosomes (and therefore confidently sequenced as diploids), 42 exhibited polymorphism at the HD genes in SNP data, indicating heterozygosity, while 20 showed no polymorphism at the HD genes. These findings confirm that heterozygosity at HD genes is no longer required for mating compatibility or completion of the life cycle. The SNP analysis further indicated a high selfing rate in these fungi: the F_IS_ mode on autosomes was at 0.83, corresponding to a selfing rate of ca s=0.91 using the formula s=2F_IS_/(1+F_IS_).

### Loss of function of HD genes in *M. scorzonerae* and genomic rearrangements

We also identified a loss of HD gene function and homozygosity of field-collected samples in a distant species, *M. scorzonerae* (Figure 1). Oxford Nanopore sequencing of two haploid genomes isolated from the same meiotic tetrad and with opposite PR alleles, revealed identical sequences for both HD1 and HD2 genes. We performed mating tests between meiotic products of 22 tetrads, which again indicated compatibility segregating a single mating-type locus: when cells were compatible for PR-determined conjugation, they also produced hyphae in 100% of the cases.

The HD1 sequences from the sequenced *M. scorzonerae* a_1_ and a_2_ haploid genomes appeared to have an uninterrupted coding sequence. The HD2 coding sequences displayed a 19 bp gap inducing a frameshift mutation in the coding region and a stop codon which would truncate the gene to 120 amino-acids instead of 487 amino-acids in *M. lagerheimii*. Using PCR amplification and Sanger sequencing, we confirmed that the frameshift is present in all the isolated cultures, as well as in the *M. scorzonerae* sample not used to produce haploid cultures. As in *M. superbum*, the analysis of 22 tetrads indicated that a single genetic factor controlled mating type in *M. scorzonerae*.

The a_1_ and a_2_ genome for *M. scorzonerae* did not assemble as well as for *M. superbum* and *M. shykoffianum.* Nevertheless, we identified two contigs in the a_1_ *M. scorzonerae* assembly that carried regions corresponding to the ancestral PR and HD chromosomes, and also parts of *M. lagerheimii* autosomes, indicating that mating-type chromosome fusion occurred, and also with ancestral autosomes (Figure S9A). The contigs carrying parts of the ancestral PR and HD chromosomes were nevertheless collinear between the a_1_ and a_2_ *M. scorzonerae* genomes, indicating ongoing recombination, with the exception of a relatively small non-recombining region around the PR locus (Figure S9B). The HD and PR genes were not in the same contigs. This suggests that there was a transient stage, as in *M. superbum* and *M. shykoffianum,* where the HD and PR genes were linked by chromosomal fusion of ancestral HD and PR chromosomes, as well as autosomes, followed by chromosomal rearrangements and subsequent loss-of-function of HD genes in mating-type determination. We could not detect telomeres however in any of these contigs.

### Progressive recombination suppression in the mating-type chromosome: evolutionary strata

We found evidence of four young evolutionary strata specific to the mating-type chromosome of *M. superbum* or of *M. shykoffianum,* therefore resulting from recent extensions of recombination cessation. Indeed, when plotting the per-gene synonymous divergence between mating-type chromosomes in *M. superbum* 1065 along the ancestral gene order (taking *M. lagerheimii* gene order as a proxy^18,41^), we found a region with significantly lower d_S_ values at the edge of the non-recombining region, while significantly different from zero (Figures 4 and S10). We called this region the “turquoise” evolutionary stratum following previous color names given to young evolutionary strata in *Microbotryum* fungi. This turquoise stratum is currently situated within the non-recombining region of the PR mating-type chromosome (Figures S4-5 and S11) while remaining still much less rearranged than the rest of the non-recombining region, as expected for a young evolutionary stratum. This very same turquoise stratum had in contrast zero d_S_ values in *M. shykoffianum* and has remained in the pseudo-autosomal region (Figures 5 and S4-S5). An additional evidence for the evolution of the turquoise stratum post-dating speciation between *M. shykoffianum* and *M. superbum* is the lack of trans-specific polymorphism (Figure 6). A single gene displayed full trans-specific polymorphism in the turquoise stratum and it was situated just at the limit with the black stratum, and it may actually belong to the black stratum. Altogether, these findings show that the turquoise stratum has moved into the non-recombining region in *M. superbum* after its divergence from *M. shykoffianum*, thus constituting a young evolutionary stratum (Figure 3).

We found footprints of another young evolutionary stratum in *M. superbum*, on the other side of the PR mating-type chromosome in the ancestral chromosomal arrangement. The d_S_ plot indeed showed high d_S_ values in the left edge of the PR chromosome (Figure 4), in the region that we called “ruby” (Figure 5).

A changepoint analysis confirmed the inference of distinct evolutionary strata in *M. superbum*, with the turquoise and ruby strata showing significantly lower mean d_S_ value than the black stratum, with discrete changes in mean values (Figure 4). The changepoint analysis even indicated that the ruby stratum was divided into two distinct sub-strata of different ages, that we called the “dark” and “light’ ruby strata, with significant different mean d_S_ values and different locations: the dark ruby stratum was rearranged, while the light ruby stratum was still at the edge of the chromosome (Figures 4 and S4). The black, turquoise, light and dark ruby strata in fact had significant different d_S_ mean values than their nearby strata (Figure S10).

The dark ruby stratum was only partly present in *M. shykoffianum* (Figure 5), suggesting either an independent origin in the two lineages or an ancestral stratum that would have partly restored recombination in *M. shykoffianum.* This latter hypothesis is supported by the lack of detected change-point in *M. superbum* within the dark ruby stratum, indicating it evolved in a single step (Figure 4). Moreover, trans-specific polymorphism was found for the genes from the right part of dark ruby stratum that have high d_S_ values in the two species (Figures 4-5).

In *M. scorzonareae,* genes around the PR locus were highly rearranged between mating-type chromosomes and compared to the *M. lagerheimii* gene order (Fig S9), indicating ancient recombination suppression, which was further supported by high d_S_ values (Fig S13). This non-recombining region extended farther than the purple and orange strata shared among all *Microbotryum* species, indicating an extension of recombination suppression specific to *M. scorzonareae.* We called this extension the emerald stratum, and it encompassed not only a part of the ancestral PR chromosome, but also a small fragment of the ancestral HD chromosome, from the middle of the MC03B *M. lagerheimii* contig: this fragment indeed displays elevated d_S_ values in *M. scorzonareae* (Fig S13) and is currently located near the PR locus because of the inter-chromosomal rearrangements (Fig S9). In addition, we detected footprints of a younger specific evolutionary stratum, called quartz, with very low d_S_ values (Fig S13), being collinear between a_1_ and a_2_ PR mating-type chromosomes but having moved from the PAR to the middle of the non-recombining region in the a_2_ PR chromosome (Fig S9). The non-zero d_S_ values extend a bit farther towards the PAR than the translocated fragment, suggesting that the quartz stratum may be larger than this translocated fragment.

We used the divergence time between M. lychnidis-dioicae and M. silenes-dioicae as a calibration point (0.42 million years^74^) to estimate the age in million years (MY) of the various evolutionary strata and their confidence intervals (CI), based on autosomal single-copy genes. We estimated the following ages: 1.55 MY for the black stratum (CI 1.21-1.92), 1.25 MY for the dark ruby stratum (CI 0.75-2.23), 0.28 MY for the light ruby stratum (CI 0.16-0.45), 1.61 MY for the emerald stratum (CI 1.03-2.33) and 0.06 MY for the quartz stratum (CI 0-0.16).

### Degeneration of the PR mating-type chromosomes

In both *M. superbum* and *M. shykoffianum*, the PR mating-type chromosome shows chaos of rearrangements between the a_1_ and a_2_ genomes and in comparison to the ancestral gene order, taking as proxy the *M. lagerheimii* PR chromosome (Figures 2, S1-S5). Similarly, expected of non-recombining regions, the PR chromosome has also accumulated transposable elements (TEs), as seen by the much higher TE content in the a_1_ and a_2_ PR chromosomes in *M. superbum* (65% and 67% of bp, respectively) compared to its autosomes (mean 35% of bp) (Figure 8). Some TE families seem to have preferentially expanded, i.e. *Copia*, Ty3 (*Gypsy)* and *Helitrons* (Figure 8). As a result, the PR chromosome has become much larger than its ancestral recombining state (Figure 8, 12,930,922 vs 942,259 bp). In contrast, the HD chromosome did not have an TE load (mean 36% of bp, Figures), in agreement with the inference of continued recombination in this chromosome.

**Figure 8:**
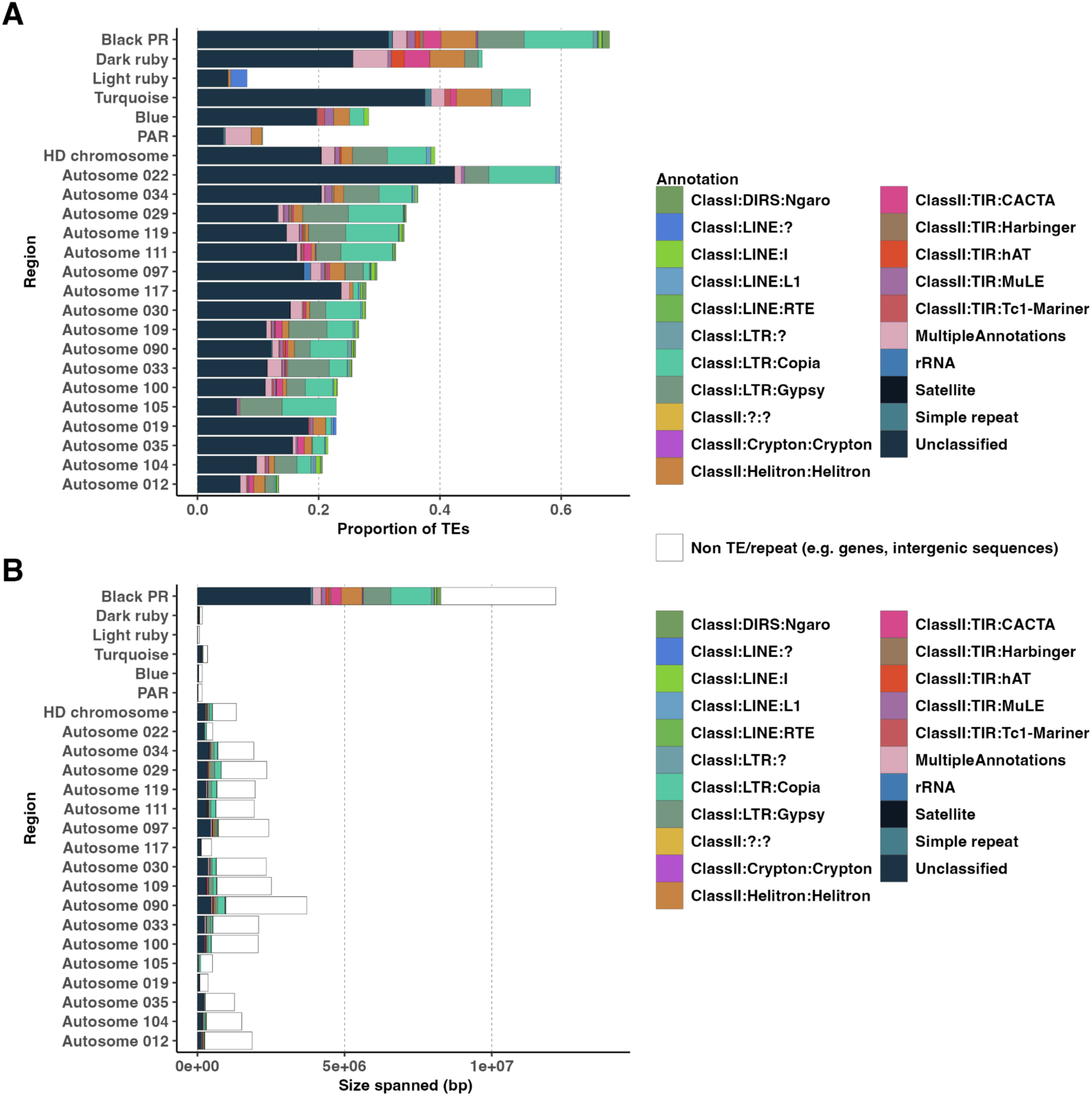
Scaffold size and transposable element (TE) content in the genome of *Microbotryum superbum* a_1_, as a proportion (A) or absolute content in base pair (bp, B). The relative proportions of transposable element classes in each scaffold is represented by the sizes of the colored bars. The PR chromosome is separated into its different evolutionary strata (blue, black, turquoise, light and dark ruby) and the pseudo-autosomal regions (PAR). The blue stratum on the HD chromosome is also separated from the rest of the chromosome. Only scaffolds longer than 300kb are displayed.

## DISCUSSION

A wide variety of breeding systems have been reported across fungi, including universal haploid compatibility, a single mating-type locus or two mating-type loci^8,14^. The role of PR and HD genes in mating-type determination and their location on different chromosomes is ancestral in basidiomycetes, corresponding to a bifactorial system, i.e., with two different mating-type loci. Having two distinct loci controlling mating compatibility can favor outcrossing, while having a single mating-type locus is advantageous under selfing: one mating-type locus yields 50% compatibility among gametes of a given diploid individual versus only 25% with two mating-type loci^56^. There have been multiple independent transitions across the Basidiomycota from bifactorial to unifactorial systems, i.e., with a single genetic locus determining mating compatibility. Unifactorial compatibility can occur through two main pathways: linkage of PR and HD loci or the loss of function at one of the mating type loci^8^. Linkage between mating-type loci has been observed in *Ustilago hordei^44^*and multiple times independently in the *Malassezia^45,46^* and *Microbotryum^18,41^* clades, as well as in other basidiomycetes^8,75-78^.

Loss-of-function at one of the mating-type loci, specifically at the PR locus, has also been observed in several basidiomycete fungi, including *Pholiota namek*o^79^, *Coprinellus disseminatus^47^*, and others^8^. To the best of our knowledge, this study in *Microbotryum* is the first known case of functional loss at the HD locus in natural populations of heterothallic basidiomycete fungi. This shows that unifactoriality, i.e. the segregation of only two mating types in progenies, can be convergently acquired through a variety of different genomic pathways in selfing fungi, for which it is beneficial. Furthermore, we found convergent events of HD gene loss-of-function after a transient HD-PR chromosome fusion, in two very distantly related anther-smut fungi. Altogether, this sheds new light on the power of selection and on the repeatability of evolution, and shows that similar phenotypes can be achieved repeatedly, via similar or different genomic mechanisms.

In addition, finding this example of an HD loss-of-function has important implications for our understanding of the sexual cycle and gene functions in basidiomycete fungi. The two tightly-linked genes of the HD locus, HD1 and HD2, are indeed thought to be essential for dikaryotic growth, as they heterodimerize to activate the dikaryotic growth and can only do so between different allelic forms at the two genes (using b_1_ and b_2_ to denote different alleles and where haploid genomes typically carry either HD1 b_1_ with HD2 b_1_ or HD1 b_2_ with HD2 b_2_, dikaryotic growth is triggered by HD1 b_1_ with HD2 b_2_ or HD1 b_2_ with HD2 b_1_). Uniting the two compatible variants in a haploid genome could allow dikaryotic growth without the need to mate with a different haplotype, as shown experimentally with a mutant carrying a chimeric homeodomain protein in the mushroom *Coprinus cinereus^49^*. The need for HD heterodimers may be bypassed for dikaryotic growth, as shown in the invasive Californian death cap mushroom, *Amanita phalloides,* that can mate as a haploid homothallic, despite the two HD genes in haploid genomes being unable to form a heterodimer^80^. Another possibility may be as in the homothallic basidiomycete yeast *Cystofilobasidium capitatum*, in which the HD1/HD2 heterodimer has likely been replaced by a HD2 homodimer^81^. This hypothesis would explain the positive selection detected on the new HD coding sequences.

Given the loss of the HD role in mating-type determination, it was intriguing to discover that the PR chromosome had incorporated a fragment of the ancestral HD chromosome and was non-recombining throughout most of its length. Indeed, the large non-recombining regions in *Microbotryum* fungi are generally partly due to the linkage of HD and PR genes, even if further extension of the non-recombining region has repeatedly evolved^18,41^. In the *Microbotryum* species parasitizing *Dianthus* and *Scorzonareae* plants analysed here, a transient stage seems to have occurred, with HD-PR linkage by fusion of the whole PR and HD ancestral chromosomes, followed by a few rearrangements and then excision of a part of the ancestral HD chromosome, likely once HD has lost its function. This hypothesis is supported by our finding of genes belonging to the ancestral HD-containing chromosome arm being now located on the PR chromosome in each of *M. superbum* and *M. shykoffianum.* Reversion to unlinked mating-type chromosomes has been suggested in *Malassezia-*related fungi^46^, but both mating-type loci kept their functions, such that reversion was likely driven by changes in mating systems. Here the chromosome fission may have followed the loss of function of the HD genes, as this loss of function would offset the benefit of HD-PR linkage. Alternatively, the fission may have occurred first, followed by the loss of function of the HD genes, selected for unifactoriality, being advantageous under selfing.

This study also provided further support for the existence of three young, species-specific evolutionary strata associated with the mating type loci in *Microbotryum* fungi. *M. superbum*, *M. shykoffianum* and *M. scorzonareae* displayed high differentiation between mating-type chromosomes and rearrangements within the non-recombining region, but lower differentiation and lower levels of gene order reshuffling in young evolutionary strata than the rest of the non-recombining region. The evolutionary cause for such stepwise and repeated recombination suppression events may be a combination of selection for less genetic load in non-recombining fragments and deleterious-mutation sheltering^31^. Indeed, if a non-recombining fragment traps fewer recessive deleterious alleles than average in the population, this fragment will have a selective advantage, but can only become fixed if it also captures a permanently heterozygous allele that will shelter its few recessive deleterious mutations when it becomes frequent enough to often be homozygous in populations^31^. The only other hypothesis that can explain such repeated evolutionary strata on fungal mating-type chromosomes is a neutral fixation of inversions or genetic differences impairing recombination^29^, but this does not explain the observed association between dikaryotic life cycle and evolutionary strata in fungi^33^. The proximal mechanism of recombination suppression may be the observed movements of fragments from the pseudoautosomal region into the non-recombining region, forming the young strata, as previously reported for the red stratum in *M. lychnidis-dioicae^17^*. However, such movements can also be a consequence, rather than a cause, of recombination suppression.

As expected, the older part of the non-recombining region on the PR mating-type chromosome shows signs of degeneration, with chaos of rearrangements and high TE load. Some TE families have preferentially expanded, i.e. *Copia*, Ty3 (*Gypsy)* and *Helitrons*, corresponding to the same families that repeatedly expanded in mating-type chromosomes of other *Microbotryum* species^69^.

Given such accumulated load that results from mutational degeneration, it has been suggested that reversion towards recombination could be selected for under the lower-loaded-sheltering hypothesis, unless dosage compensation evolves^32^. Our scenario actually proposes an early reversion of recombination suppression along a part of the ancestral HD chromosome. The chaos of rearrangements on both mating-type chromosomes in the oldest evolutionary strata, however, reinforces the view that rearrangements accumulate following recombination suppression such that it seems challenging to restore recombination when load has accumulated^31,33^. Recombination restoration would be even more difficult if the proximal mechanism of recombination suppression is the movement of pseudo-autosomal regions into non-recombining regions, as observed for young evolutionary strata. Recombination restoration would indeed then require the exact same movement back, with exactly the same breakpoints, which seems highly implausible. Our findings suggest that recombination restoration may occur when rearrangements are not too extensive, as evidenced by i) the footprints of recombination suppression followed by chromosome separation and recombination restoration in the HD chromosome arm, ii) the reversion of a part of the dark ruby stratum to a recombining state in *M. shykoffianum.* However, recombination restoration now seems impossible in the non-recombining region of the PR chromosome given the chaos of rearrangements there in both mating-type chromosomes.

In conclusion, our study brings novel insights into the evolution of breeding systems and sex-related chromosomes, reporting unprecedented cases of loss-of-function of HD mating-type genes in fungi, and fusion of ancestral mating-type chromosomes without linkage of mating-type genes. Our findings further reinforce the view that recombination suppression can extend progressively in organisms without sexual antagonism.

## Material and methods

### Study systems

*Microbotryum* fungi, parasitizing *Dianthus* and *Silene* species, castrate plants by producing their spores in the anthers of host plants and aborting the ovaries. Diseased plants are identifiable by dark-colored spore masses in their flowers. Most *Microbotryum* fungi are specialized on a single host species and most host species harbor a single *Microbotryum* species, especially in the *Silene* genus^51,52,53, 82^. One exception is the clade of *Microbotryum* fungi parasitizing wild carnation relatives in the pink family (*Dianthus* genus, Caryophyllaceae), in which several closely-related and sympatric *Microbotryum* species parasitize multiple *Dianthus* plant species, with host range overlapping to some extent^52,53, 83, 84^ (Figure 1). Three species have been formally described so far in this group^84^, but they are challenging to recognize based on their host plant or morphology. We therefore assigned strains to species based on the published sequences of the gene barcodes used for describing the species (EF1-alpha and beta-tubulin;^52,84^; see below). *Microbotryum scorzonerae* parasitizes flowers of *Scorzonera humilis*, in the Asteraceae family^85^. Unlike other *Microbotryum*, *M. scorzonerae* does not cause spore formation on anthers alone. It instead causes spore formation within the entire floral head (Figure 1).

### Genomic analyses

#### Strains, DNA extraction and species identification

We extracted DNA as previously described^17,18^. For the *Dianthus* strains, DNA sequencing based on PacBio (Pacific Bioscience) HiFi long-read sequencing was performed at the GenoToul sequencing facility (Toulouse, INRAE, France) in 2021. We isolated haploid sporidia of opposite mating types from a single meiosis (tetrad) from the strain 1065, collected in July 2011 on *Dianthus pavonius*, in Italy (coordinates 44.189, 7.688, near the Garelli Refugium in the Alps) by Michael Hood, Janis Antonovics and Emme Bruns. Based on preliminary analyses of SNPs, we also identified a strain belonging to another *Microbotryum* species parasitizing *Dianthus* plants: strain C212 (CZ_D24), collected in 2016 on *Dianthus carthusianorum*, in the Czech Republic, coordinates 49.639759, 14.201153 (Bohemia, The Vltava river basin, to the north of Zduchovice) by Klara Koupilova. This later strain had been cultivated on Petri dishes at the haploid stage after spreading diploid teliospores that each underwent meiosis; we therefore sequenced a mixture of haploid sporidia resulting from multiple meioses from a single diploid individual. The mixture is thus equivalent to a diploid genome. Cultivation of sporidia was performed on PDA (potato dextrose agar)^86^. The samples were collected in Italy, which is not a party of the Nagoya protocol, and in the Czech Republic, which does not regulate access to its genetic resources in relation to the Nagoya Protocol. Using the best BLAST hits of the EF1-alpha and beta-tubulin genes against the reference sequences^52,84^, we assigned the strain 1065 to *M. superbum* (aka MvDsp2) and the strain C212 to *M. shykoffianum* (aka MvDsp1)^52,84^. For the *M. scorzonarae* strain, obtained in 2018 from Cefn Cribwr, Wales, by Julian Woodman, sporidia of opposite mating types were isolated from a single meiosis (tetrad). Although a party to the Nagoya Protocol, the UK does not regulate access to its genetic resources. From cultures on PDA, extracted haploid genomic DNA was used to generate long-read sequences with MinION (Oxford Nanopore Technologies), and short-read Illumina sequencing.

#### RNAseq from M. superbum and M. intermedium

We generated RNAseq data for gene prediction and HD gene expression analysis. For optimizing gene prediction, we sequenced RNA from one of the focal species, *M. superbum,* and also from an outgroup to our studied *Microbotryum* species, *M. intermedium*. For *M. intermedium,* we used the strain BM12-12.1 (the same strain as used previously for genome sequencing^17,39^), under two different conditions: 1) haploid cells of a single PR mating type (a_1_ or a_2_) grown on nutrient medium (PDA) and then mixed in equal proportions of the two mating types before RNA extraction (“H” condition), as well as additional replicates grown separately on nutrient agar, whose transcriptomes were analyzed separately, and 2) mixtures of cells of the two mating types under mating conditions (“M” condition), i.e. on water agar. Total RNA was isolated from haploid *M. intermedium* cells using the Monarch® Total RNA Miniprep Kit (New England Biolabs). For the nutrient medium condition, haploid strains (a_1_ or a_2_) were streaked onto potato dextrose agar (PDA) and grown for 3-4 days at 22 °C. Cells were scraped, suspended in DNA/RNA protection solution, disrupted mechanically in the presence of acid-washed glass beads (0.5 mm dia), and then total RNA was extracted following the manufacturer’s protocol. For the mating condition, haploid a_1_ and a_2_ cultures were first grown separately in yeast extract peptone dextrose (YPD) broth overnight. Cell density was measured with a spectrophotometer. The concentration of each culture was adjusted to O.D.600 1.0. Equal volumes of the two haploid cultures were mixed and plated on water agar. These plates were incubated for four days at 14°C, after which wet mounts were prepared from each “mated” plate to verify the presence of conjugation tubes, indicating active mating behavior. Cells were scraped and then total RNA was isolated using the Qiagen RNeasy Plant Mini Kit. After total RNA isolation, several quality control measures were taken. Concentration and purity were assessed using a NanoDrop 2000 spectrophotometer (Thermo Scientific); 260/280 and 260/230 ratios>1.8 were considered satisfactory for RNAseq application. Additionally, cDNA was prepared and used as a template for intron-spanning primers in PCR reactions to verify the lack of genomic DNA contamination. Bioanalyzer analysis was completed to detect intact 18S and 23S RNA as a measure of overall RNA quality. After passing all quality control measurements, RNA was sent for sequencing to CD Genomics (Shirley, NY, USA). Three replicates were sequenced per condition and means across the three replicates were analyzed.

For *M. superbum,* we used another strain from the same *D. pavonius* population as the strain 1065 (coordinates 44.191, 7.685, near the Garelli Refugium in the Italian Alps) for RNASeq, a_1_ and a_2_ haploid strains were used separately as well as pooled in mating experiments. For extractions from haploid cells grown on PDA, we used the Zymo -----RNA Extraction Kit (Zymo Research, Irvine, California). Prior to RNA extractions, each haploid fungal strain was grown on two PDA Petri dishes at 28°C for 24 hours. Extractions were performed on fungal cells from the two PDA Petri dishes for each haploid sample replicate. Fungal mating was accomplished in the following steps. Haploid *M. superbum* a_1_ and a_2_ cells were grown on YPD agar for four days prior to the mating assay. Haploid cells were then suspended in distilled and autoclaved water before adjusting to 10^9^ cells/mL. Suspended fungal cells were combined in equal proportions before being spotted in 50 μl spots on water agar Petri dishes. Plates were left to incubate at 13° Celsius for 48 hours. Resulting cells were visualized under a light microscope to confirm the occurrence of mating via the formation of conjugation tubes between cells. Under these conditions we typically observed about 30% of the cells engaged in mating. Extractions using the Zymo RNA extraction kit were performed on mated fungal tissue that was aggregated from the water agar petri dishes for each mated sample. RNAseq of three biological replicates per condition (mated, haploid a_1_ and haploid a_2_) were performed by CD Genomics (Shirley, New York).

Three *D. pavonius* plants infected with *M. superbum* were obtained from Emme Bruns at the University of Maryland. For generating these diseased plants, seeds of *D. pavonius* were collected from two populations in Aug 2016 in Italy (Duca coord. 44.1946327, 7.66067016 and Garelli coord. 44.189128, 7.688879), which is not a party of the Nagoya protocol. They were reared in greenhouse conditions planted in Sept 2016, reared and crossed within populations in 2017 to produce seeds at the University of Maryland. F1 seeds were planted and inoculated in 2018 with a strain from the Duca population, and scored as diseased 2020. The plants (DE 10B 3; HL-2 duca 1061; De 10A 8) were used for RNAseq production. Four different flower bud size binspools were used in these analyses: 11-12 mm, 13-14 mm, 15-16 mm, and 18-20 mm. These represented different stages of both host and parasite development. RNA extractions from plant tissue were performed using Qiagen -----RNA Extraction Kit. Flower buds were harvested from plants that were maintained in a controlled growth chamber. The number of flower buds used for each extraction was dependent on the size of the buds that were collected. All extractions with flower buds that fell into the 11-12 mm bin category were completed with at least two flower buds while extractions on buds in the 13-14 mm or larger bin categories used one flower bud per reaction. In total we sent 9 samples from three plants for RNAseq (CD Genomics).

#### Expression of HD genes

For the gene expression analysis on HD genes, we used only the conditions under which we had multiple technical replicates and for which HD genes are expected to be expressed, i.e, the three 11-12 mm *M. superbum* replicates obtained from the same plant and the three replicates from each of the culture conditions (mated at 48 h and haploid cultures). Expression of HD genes in *M. lychnidis-dioicae* and *M. intermedium* M. lychnidis-dioicae, or PDA, M. intermedium, and for both under mating conditions at 48 h^39,76^ was assessed as a control of expectations for functional HD genes, in two or three replicates for each of three culture conditions (haploid cultures of alternative mating types grown on water, *M. lychnidis-dioicae*, or PDA, *M. intermedium*, and for both under mating conditions at 48 h^39,87^) and from infected flowers at a late stage^88^, see Table S3. To check whether HD genes were expressed, reads were mapped with minimap2 2.26-r1175^89^ with the -ax sr option, depth per base extracted with samtools depth function^90^, explored with jbrowse 2^91^ and visualized with plotkaryotypeR^92^.

#### Genome assemblies and gene prediction

For *M. superbum* 1065, we sequenced separately two haploid genomes, *a priori* of compatible mating types based on their position in the isolated linear tetrad. We assembled these two genomes separately with Canu 2.2^93^ using the fastq reads files as input, and setting the -pacbio-hifi parameter. For the strain collected on *D. carthusianorum*, we sequenced sporidial cultures expected to correspond to mixtures of meiotic products from a diploid individual. Therefore, we used the software Hifiasm 0.16.1-r375^94^, which can separate haplotypes, with default parameters. We performed quality assessment of the resulting assembly by computing N50 and L50 statistics and by assessing genome completeness with BUSCO v5.4^95^ using the basidiomycota_odb10 as reference genes.

The two alternative mating types of M. scorzonarae were sequenced on Amherst laboratory ONT platform. Longer than 1000bp of raw ONT reads were kept for genome assemblies. Trimmed reads were assembled with Canu v1.7b with default parameters ^93^. The resulting assemblies were polished with Pilon v1.22 (https://github.com/broadinstitute/pilon/wiki) with default parameters, using Illumina 150 bp paired-end reads belonging to the two mating types of the same strain. To that end, short-read sequences were quality assessed using FastQC (https://www.bioinformatics.babraham.ac.uk/projects/fastqc/), trimmed with Trimmomatic () using default parameters. Then 3 rounds of polishing were run iteratively with Pilon, with Illumina short reads aligned to the polished assemblies obtained from the previous round and indexed at each iteration using bwa-mem2 ^96^. We then assessed Genome completeness with BUSCO v5.4 ^95^ using the basidiomycota_odb10 as reference genes.

Repeated regions and low complexity DNA sequences were identified using RepeatModeler^97^. The genome was then softmasked using RepeatMasker open-4.0.9^98^ with the *de novo* detected repeats as well as custom libraries of candidate transposable elements (TEs) from our previous studies^41^ and existing fungal TEs from RepBase^99^. To minimize the rates of false negatives in the gene prediction steps, we excluded from the softmasking TEs that were labeled as “unknown” as their TE status was uncertain.

Genes were then predicted on the soft-masked genome using the software Braker version 2.1.6^100^ which uses Diamond, ProtHint 2.6.0, GeneMark 4 and AUGUSTUS 3.4.0. To help gene prediction, we combined RNAseq data from *M. superbum*, *M. intermedium* and *M. lychnidis-dioicae*, and a set of highly conserved protein sequences previously identified across multiple *Microbotryum* species. This set of proteins contains fully conserved single-copy genes not overlapping with the mating-type chromosomes from 18 published *Microbotryum* genomes, all predicted with the same pipeline^17,18,41,65^. The Illumina RNAseq data from 12 *M. superbum* and 6 *M. intermedium* samples generated here together with 5 *M. intermedium*^39^ and 17 *M. lychnidis-dioicae*^87^ samples previously obtained (Table S3) were processed with fastp^101^, aligned to the reference genome with the software GSNAP and GMAP^102^, combined and sorted with samtools^90^ and filtered with Augustus^103^ to keep only the unique best match (--uniq option). Following Braker’s documentation, we performed gene predictions separately for RNAseq data and protein databases. For the gene prediction, we ran five rounds of gene prediction and kept the run showing the highest BUSCO score. We eventually combined the protein and RNAseq gene predictions using TSEBRA^104^ and assessed the quality of the final gene set with BUSCO using the basidiomycota_odb10 as reference using a dedicated workflow (https://github.com/QuentinRougemont/genome_annotation). D*e novo* detection of transposable elements (TEs) was performed as described previously^69^.

#### Orthogroup construction

We performed orthology reconstruction using a set of 47 published genomes from our previous work^39,41^ and including the newly assembled genomes for three species studied here. The *Rhodosporidium babjavae* genome was used as an outgroup^41^. We identified transposable elements and predicted genes in each genome using Braker v2.1.6 with the protein database built as explained above and we used in addition, for the *M. superbum*, *M. intermedium* and *M. lychnidis-dioicae* genomes, RNAseq data from these species (n = 12, 11 and 17 conditions for *M. superbum*, *M. intermedium* and *M. lychnidis-dioicae* respectively, Table S3). When multiple isoforms were present, we kept the gene prediction with the longest transcript. This extensive dataset was then fed to OrthoFinder^105^ with default parameters to reconstruct orthogroups.

#### Identification of mating-type chromosomes

We identified the contigs belonging to the mating-type chromosomes by i) detecting the contigs carrying PR and HD genes using BLASTN 2.6.0^106^ with the gene sequences from *M. lagerheimii* as queries^72, 107^, ii) plotting the ortholog links for all contigs of our focal species against the reference genome of the *M. lagerheimii* strain 1512 (a_1_)^17^ using Rideogram^108^ iii) plotting the ortholog links using Rideogram between the a_1_ and a_2_ genomes of our focal species.

We also generated circos plots using circlize^109^. In circos plots, we oriented contigs in the direction involving the fewest inversions and keeping the recombining regions of the mating-type chromosomes, called pseudo-autosomal regions (PARs), at the edges and non-inverted between mating-type chromosomes.

#### Analysis of selection on HD genes

We aligned the coding sequences of the HD2 gene for all the *Microbotryum* species with available genomes (Figure 1) using the software MACSE v2^110^. The HD2 gene that was predicted as two separate genes in the a_2_ genome of *M. superbum* was manually merged to form one sequence, conserving the alignment with the other species. We used the RELAX method from the hyphy package^111^ to detect relaxed or intensified selection on the branch of the HD2 gene in the a_2_ genome of *M. superbum*, compared to the HD2 gene sequence in the a_1_ and a_2_ *M. lagerheimii* branches.

#### Detection of evolutionary strata

We used as a proxy of the ancestral state the chromosomal arrangement and gene order of *M. lagerheimii,* as done in previous studies^17,65^. In *M. lagerheimii,* PR and HD loci are located on different chromosomes^17^. For each species, synonymous divergence (d_S_) was computed from the alignment of a_1_ and a_2_ allele sequences, obtained with macse^112^, using the yn00 v4.9f program of the PAML package^113^ and plotted using the ggplot2 library of R^114^. Pseudo-autosomal, recombining regions were identified as regions with null synonymous divergence on d_S_ plots, as expected in these highly selfing fungi, and with the same gene order between mating types and as in *M. lagerheimii,* as assessed on circos plots. Non-recombining regions were identified in contrast as regions with non-null synonymous divergence on d_S_ plots, and being most often rearranged compared to the ancestral gene order in synteny plots. In order to objectively identify the limits of evolutionary strata, we looked for changes in mean d_S_ along the ancestral gene order along the mating-type chromosomes using a changepoint analysis performed using the R package mcp^115^. This analysis relies on Bayesian regression to infer the location of changes in means of the d_s_ values.

#### Telomeres and centromeres

To detect telomeres, we counted the occurrence of the telomere-specific motif ‘TTAGGG’ and its reverse complement ‘CCCTAA’ within 1kb windows along the contigs. Peaks occur when telomeres are present as they are constituted by multiple repeats of this motif. Centromeres were identified by searching for the centromeric repeats previously described in *Microbotryum* fungi^41,42^.

#### Trans-specific polymorphism

Trans-specific polymorphism (i.e., the clustering of alleles per mating type across species) can be used to study the age of recombination suppression linking genes to mating-type loci. Indeed, as soon as a gene is fully linked to a mating-type locus, its alternative alleles will remain associated to alternative mating types, even across speciation events. Mutations will therefore accumulate independently in alleles associated with alternative mating types. In a genealogy, alleles will therefore be grouped according to mating type rather than according to species. The node at which the alleles associated with the alternative mating types diverge indicates the time of recombination cessation^17,18,23^.

We performed codon-based alignment with macse v2.05^110^ of one-to-one orthologs of genes ancestrally located on mating type chromosomes (416 genes). Gene trees were obtained with IQ-TREE 1.6.1^116^ with default parameters. Tree topologies were assessed using a homemade awk command line, available on demand. Trees were separated into three categories: species tree (a_1_ and a_2_ are grouped together within the species node), full trans-specific polymorphism (both a_1_ are grouped together and both a_2_ are grouped together) and partial trans-specific polymorphism (when only the a_1_ or the a_2_ are grouped together, or some alleles are missing).

A large part of the HD chromosome is collapsed in the *M. shykoffianum* genome assembly, which means that it was not correctly separated between a_1_ and a_2_ genomes during the assembly. Therefore, we could not run trans-specific polymorphism analyses in this region because the genes could only be predicted in the a_1_ genome assembly. However, the assembly collapse means that the a_1_ and a_2_ alleles were highly similar, with therefore likely no trans-specific polymorphism.

#### Dating of evolutionary strata

For each gene assigned to an evolutionary stratum in either of the three focal species, we identified the orthologous genes in the mating-type chromosomes of four reference species (eight haploid assemblies): *M. lagerheimii* and *M. intermedium*, displaying an ancestral-like gene order ^17^, and *M. lychnidis-dioicae* and *M. silenes-dioicae*, two species for which the divergence time has been estimated ^74^. We filtered out orthologous groups with a copy number different from one in any of the haploid genomes. For the dark ruby stratum we only kept the genes assigned to the stratum in both *M. shykoffianum* and *M. superbum*. We identified a minimum of 6 (dark ruby stratum) and a maximum of 31 (black stratum) orthologous groups per stratum. We obtained codon-based alignments for each orthologous group with translatorX ^117^ using muscle ^118^ as protein aligner. For each stratum, we concatenated the codon-based alignments of its orthologous groups (alignment length range: 9528 - 75651 base-pairs) and reconstructed an ultrametric tree with the least square dating method ^119^ under the TPM+F+G4 substitution model, as implemented in iqtree2 (version 2.2.6 ^116^). We used the divergence time between *M. lychnidis-dioicae* and *M. silenes-dioicae* as a calibration point (0.42 million years^74^).

#### Illumina sequencing and analysis

A total of 165 samples were collected on several diseased *Dianthus* plants from various species (*D. pavonius, D. seguieri, D. carthusianorum, D. hyssopifolius, D. albinos, D. superbus, D. caryophyllus, D. deltoides* and *Stellaria holostea*) in several locations across Europe. For each strain, diploid spores from anthers of a single flower were spread on a Petri dish containing potato dextrose agar (PDA) and ampicillin, and let grow at 23°C and under artificial light. Spores in a given flower are produced by a single diploid individual^120^. When placed on a nutritive media, the diploid spores undergo meiosis and the resulting haploid sporidia then replicate clonally. There were thousands of haploid sporidia harvested on the Petri dishes representing the numerous meiotic products of a single diploid individual. However, lethal alleles are sometimes found linked to one mating type in populations, so that some mixtures of recovered sporidia can include only the a_1_ or a_2_ mating-type chromosome^58,66^. Cells were harvested from the medium and stored at -20°C until DNA extraction, using the Macherey-Nagel NucleoSpin Soil® kit following the manufacturer’s instructions. The quality of the DNA extracted was then assessed by measuring the ratio of 230/260 and 280/260 nm with a NanoDrop 2000 spectrophotometer (Thermo Scientific). A Qubit 2.0 fluorometer was used to measure DNA concentration. Preparation of DNA libraries for sequencing was performed using the Illumina TruSeq Kits®. Paired-end libraries of 2*150 bp fragments with an insert size of 300 bp were prepared with Illumina TruSeq Nano DNA Library Prep® kit. Sequencing was performed on a HiSeq 2500 Illumina sequencer, with an average coverage of 10-15X. We sequenced 49 genomes of *Microbotryum* strains from *D. carthusianorum*, 87 from *D. pavonius*, 12 from *D. seguieri*, and 17 from other hosts across western Europe. The samples were collected in Italy, which is not a party of the Nagoya protocol, in the Czech Republic, Germany, Finland, UK and the Netherlands, which do not regulate access to its genetic resources in relation to the Nagoya Protocol, in Switzerland, where we just need to keep track of information for fundamental research, and in France, where there was an exception for micro-organisms at the time of sampling.

We followed GATK pipeline best practices for read mapping and SNP calling. Illumina reads from the 165 whole genomes were mapped onto the *M. superbum* reference genome. We constructed a “hybrid” reference genome by merging the whole genome MvDp−1065−A1 and the two mating-type chromosomes of the MvDp−1065−A2 genome (MvDp−1065−A2_tig00000068 and MvDp−1065−A2_tig00000031, respectively harboring the a_2_ PR and HD alleles). We mapped the reads against the hybrid reference genome using BWA-MEM2^121^ (https://github.com/bwa-mem2/bwa-mem2, t-4 option). For mating type-chromosomes, only reads mapping to a single location were considered for further analyses. We used GATK Haplotype Caller to obtain gVCF from the bam mapping files using the haploid flag (ploidy =1) for the mating-type chromosomes, and using the diploid flag (ploidy =2) for the rest of the genome. We used CombineGVCFs to combine gvcf of the autosomes and mating-type chromosomes for a given individual, and to obtain the vcf file. We filtered out highly genetically related and duplicate individuals (i.e., individuals with pairwise KING-robust kinship estimates among individuals >0.354)^122^ with plink version 2.0 ^123^, and individuals with more than 20% of positions with missing genotype. We computed pairwise linkage disequilibrium between genome-wide biallelic sites with minor allele frequency higher than 0.25 using Plink v1.90b6.26. We used for this analysis only reads mapped to the genome MvDp−1065−A1 and the 149 strains that passed filtering.

In our analysis of homozygosity of HD genes, we first checked the ploidy of the sequenced samples at mating-type chromosomes, as they were mixtures of meiotic products with possibly haplolethal alleles associated with mating types. For this goal, we performed a remapping of the reads on the hybrid reference genome. Two autosomes were used as controls: tig00000012 and tig00000097. We calculated the median coverage along the two autosomes, as well as along the PR and HD chromosomes. Strains with a median coverage lower than six reads for the control autosomes were not considered further. The median coverage of the HD and PR chromosomes were divided by the median coverage of the control autosome (tig00000012), to get their relative coverage. Strains with a relative coverage higher than 0.7 for PR a_1_ or higher than 0.8 for PR a_2_ were considered as haploid (23 a_1_ and 14 a_2_ strains). Strains with a relative coverage around 0.5 (between 0.35 and 0.7) for PR a_2_ also showed a coverage around 0.5 for PR a_1_ and were therefore considered reliable diploid (62 strains). The 17 other strains were considered as having “unknown ploidy”. The results were consistent when using the other autosome (tig00000097) as a control. Heterozygosity of the HD chromosomes was calculated for each strain on the vcf files using the command -het of vcftools, and was defined as the observed number of heterozygous sites divided by the total number of sites.

#### Tetrad isolation and PCR analyses

We pulled out tetrads of *M. scorzonerae* from *Scorzonera humilis* (strain 1528, collected in 2018 in Cefn Cribwr, Wales, UK 51°32’08”N 3°38’52”W by Julian Woodman), *Tragopogon pratensis* (strain 1535 collected in 2018 in Länsimäentie, Helsinki, Finland, 60°13’54.753"N 25°6’57.1716"E, by Juha Tuomola), *Tragopogon pratensis* (strain 1544, collected in 2018 in De Weere, Netherlands, 52°44’13.33"N 4°59’22.5"E, by Justus and Maureen Houthuesen-Hulshoff), and *Tragopogon porrifolius* (strain 1560 collected in 2018 in De Weere, Netherlands, 52°44’13.33"N 4°59’22.5”E 2018, by Justus and Maureen Houthuesen-Hulshoff). The samples were collected in Finland, the Netherlands and the UK, which do not regulate access to their genetic resources in relation to the Nagoya Protocol. We also pulled out 20 tetrads from seven *M. superbum* strains collected on *D. pavonius* in 2019 in Garelli (Italy 44.1887, 7.6876). Italy is not a party of the Nagoya protocol.

To obtain pure haploid cultures from *M. scorzonerae* and *M. superbum* samples, teliospores were first germinated by plating on water agar and incubated at room temperature for approximately 12 hours. In *Microbotryum* fungi, a linear basidium containing a meiotic tetrad of cells forms during germination^58^. Meiosis occurs in the basidium, yielding four haploid cells. The four haploid basidium cells then undergo mitosis to form haploid sporidia. These sporidia can be separated from the basidium and isolated into separate cultures by micromanipulation^58^. In order to assess segregation patterns of the mating-type loci across multiple meioses, sporidia cells were isolated from 22 tetrads in *M. scorzonerae* and 20 tetrads in *M. superbum* samples from seven *M. superbum* strains collected from *D. pavonius*. Each sporidium cell was then moved by microcapillary pipetting to a potato dextrose agar (PDA) plate, and incubated at room temperature. Two of every four cells in each tetrad divided only a few times after isolation in *M. scorzonerae* and then stopped division. This inviability has been seen in other *Microbotryum* species and is called haplo-lethality^58^. Sporidial cultures which grew successfully were maintained continually on PDA and were stored as samples in silica powder at -18°C.

Primers were designed, using Primer3^124^, that would specifically amplify, for each the PR, HD1 and HD2 mating-type genes, either of the two alleles that were present in the two haploid reference genomes of *M. superbum*, *a priori* compatible for mating given their opposite position in the tetrad^58^ (Table S1).

For amplifying fragments of PR or HD genes, DNA templates were prepared by suspending portions of fungal cultures (quantity not exceeding the tip of a sterile toothpick) in 40 μL of ultrapure water. Boiling PCR was performed in a final volume of 30 μL containing 3 μL of fungal suspension, 15 μL of 2X Buffer (containing a mix of dNTPs 5mM in addition of bovine serum albumin (BSA) 20mg/ml, W1 detergent 1%, home made 10X Buffer (Ammonium Sulfat 1M, Tris HCl 1M, MgCl_2_6H_2_O 1M) and sterile deionized H_2_O), 1.2 μL of each forward and reverse primers (10 μM), 0.15 μL of Taq DNA Polymerase (5 U/μL, MP Biomedicals) and 9.8 μL of sterile H_2_O. The touchdown PCR amplification protocol consisted of 5 min at 94°C for pre-denaturation, 10 cycles of denaturation at 94°C for 30 sec, annealing at 60°C (with 1°C decrements from 60°C to 50°C at every cycle) for 30 sec, and extension at 72°C for 1 min. This was followed by a further 30 cycles of denaturation at 94°C for 30 sec, annealing at 50°C for 30 sec, and extension at 72°C for 1 min. The reaction was finished with a final extension for 5 min at 72°C. The PCR products were revealed and confirmed by the GelRed Nucleic Acid Stain 10000X (Biotium) and photographed with UV transillumination using the VWR Genosmart gel documentation system after loading on 1.5% agarose (Nippon Genetics Europe GmbH)-TAE gels. PCR products were sequenced using Sanger technology at Genewiz.

#### Mating tests and plant inoculations

To check for compatibility between cultures in tetrads, mating tests were performed between pairs of cultures from the same meiotic tetrad. Six-day old cultures were suspended in sterile water. Suspensions were then combined in equal amounts and plated in small spots, less than 10μl each, onto water agar plates with 0.5ml phytol solution and onto water agar plates with 0.5 ml sterile water. The phytol solution was 0.006% phytol and 0.8% ethanol^58^. Phytol has been shown to induce hyphal growth in *Microbotryum* fungi^125^. Plates were refrigerated at 5°C. Spots were checked for hyphae using an inverted scope between 2 and 4 days after plating. We used cultures of *M. superbum* sporidia of opposite PR mating types but identical HD alleles to inoculate seeds of *Dianthus chinesis* as previously described^58^. We grew the plants in the greenhouse and checked for symptoms in anthers. The species *D. chinesis* was used as it is easier to grow under the greenhouse and make it flower than *D. pavonius* or *D. carthusianorum* and the focal *Microbotryum* fungi are found on various Dianthus hosts in nature.

## Softwares for figures

For figures illustrating synteny, we used the package circlize^126^ or Rideogram^108^ based on alignments performed with mummer^127^. We used ggplot2 for plotting d_S_ values. For RNAseq mapping illustration, we used the package karyoploteR^92^.

## Data availability

RNASeq data for all experiments listed in this section are available in GEO at NCBI, Accession Numbers/Record GSE233508 for the *M. superbum* data.

## Acknowledgements

This work was supported by the Louis D. Foundation award and EvolSexChrom ERC advanced grant #832352 to T. G, a National Institute of Health (NIH) grant R15GM119092 to M. E. H., and a National Science Foundation (NSF) grant #2007449 to M.H.P. We thank Klara Koupilova, Janis Antonovics, Emme Bruns and Juha Tuomola for the *M. v. Dianthus* strains, Justus & Maureen Houthuesen-Hulshoff, Julian Woodman, Juha Tuomola for the *M. scorzonerae* strains, and Emme Bruns for *Dianthus* plants. We thank Julian Woodman for the picture of *M. scorzonareae*.

## Author contributions

T.G., M.E.H., A.C. and R.R.d.l.V. conceptualized the study, acquired funding and supervised the study. M.E.H., T.G. and A.C. acquired the *Microbotryum* strains. A.S. extracted DNA for PacBio sequencing. M.E.H. and W.J.M. sequenced and analysed the *M. scorzonerae* genome. E.L., P.J., Q.R., L.B. and R.R.d.l.V. analyzed the PacBio genomes. A.C. acquired the Illumina genomes and performed SNP calling. P.J. and E.L. analysed the Illumina genomes. M.E.H. and J.G. performed tetrad isolation and mating tests. A.L., E.C., M.E.H. and J.G. performed tetrad segregation analyses. M.H.P. and R.K.H. acquired the expression data. R.R.d.l.V. analyzed the RNAseq data. E.L., P.J. and R.R.d.l.V. produced the figures. T.G. and E.L. wrote the manuscript draft with contributions by P.J., M.E.H., Q.R., M.H.P. and R.K.H. All authors edited the manuscript. The final version of the manuscript was prepared by E.L., R.R.d.l.V. and T.G.

## SUPPLEMENTARY FIGURES

**Figure S1:**
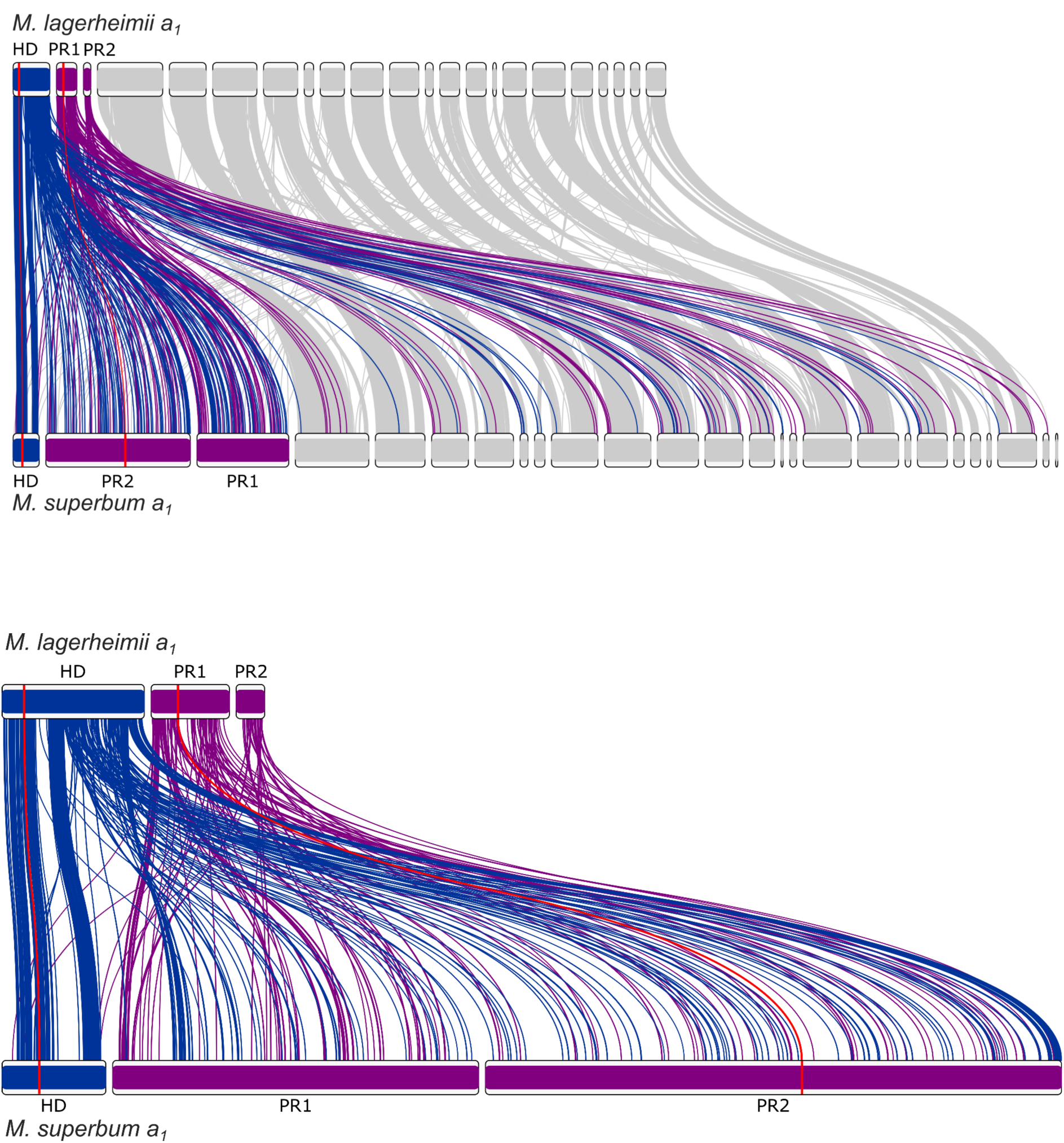
Synteny and rearrangements between the a_1_ genomes of *Microbotryum superbum* and *M. lagerheimii*, with all contigs (top) or only the mating-type chromosomes (bottom). The *M. lagerheimii* genome represents a proxy for the ancestral state before recombination suppression and with two separate mating-type chromosomes. The HD mating-type chromosome from *M. lagerheimii* is represented in blue and the PR mating-type chromosome in purple (splitted into two scaffolds), with links to orthologous genes to the *M. superbum* mating-type chromosomes. The positions of the HD and PR mating-type genes are indicated in red and by red links between the two genomes. The PR chromosome shows a substantial increase in size and a chaos of rearrangements.

**Figure S2:**
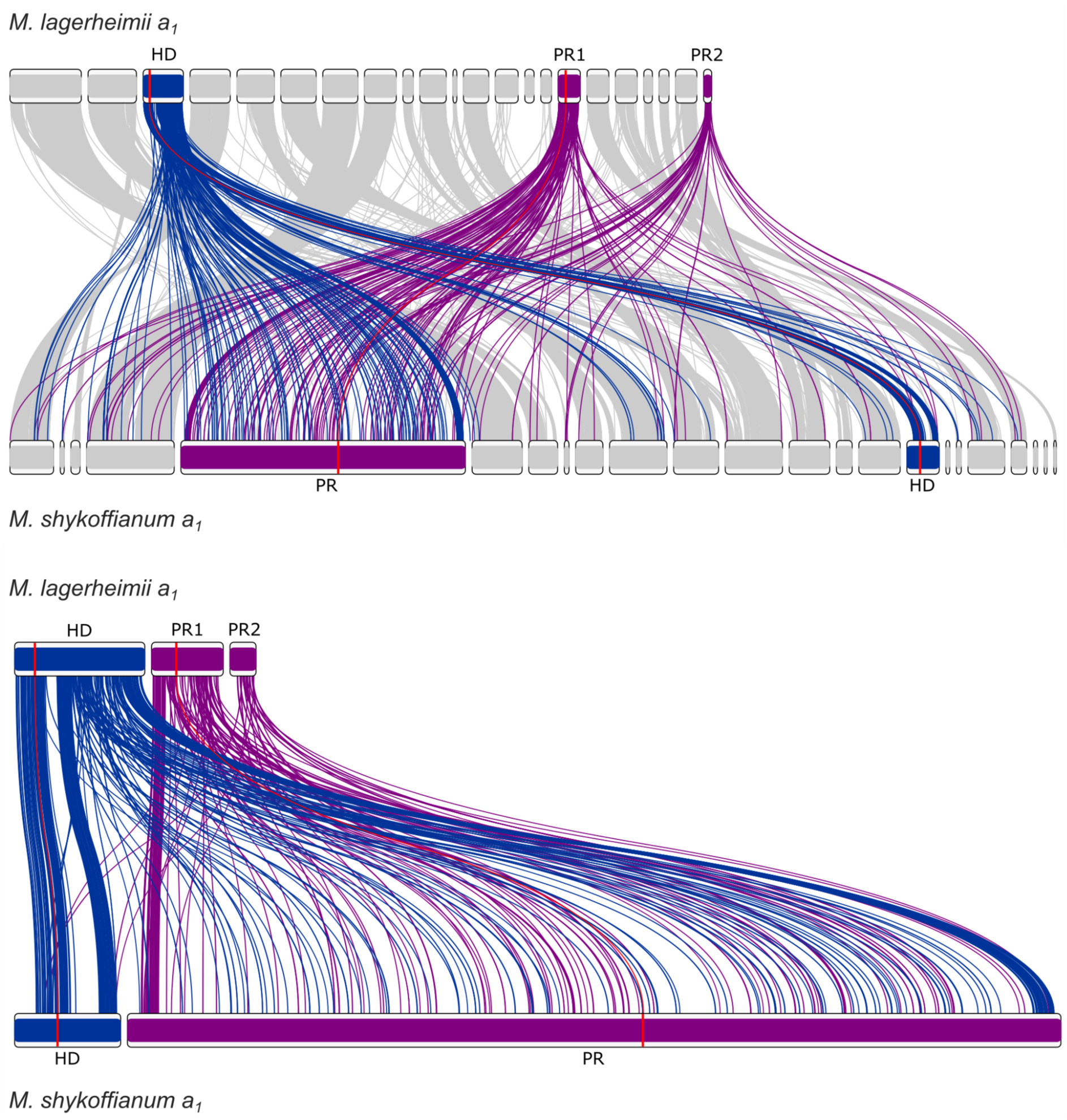
Synteny and rearrangements between the a_1_ genomes of *Microbotryum shykoffianum* and *M. lagerheimii*, with all contigs (top) or only the mating-type chromosomes (bottom). The *M. lagerheimii* genome represents a proxy for the ancestral state before recombination suppression and with two separate mating-type chromosomes. The HD mating-type chromosome from *M. lagerheimii* is represented in blue and the PR mating-type chromosome in purple (splitted into two scaffolds), with links to orthologous genes to the *M. superbum* mating-type chromosomes. The positions of the HD and PR mating-type genes are indicated in red and by red links between the two genomes. The PR chromosome shows a substantial increase in size and a chaos of rearrangements.

**Figure S3:**
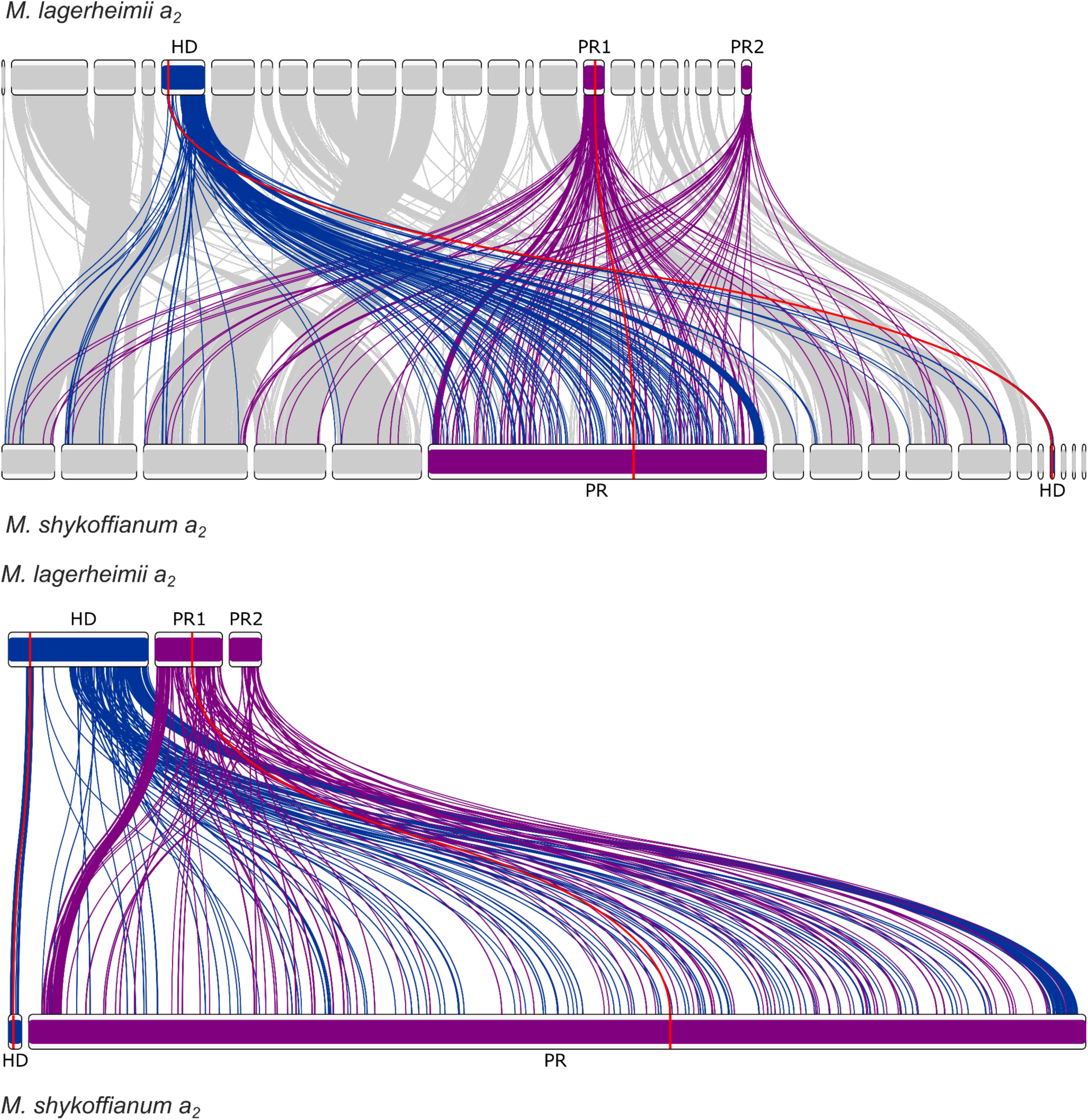
Synteny and rearrangements between the a_2_ genomes of *Microbotryum shykoffianum* and *M. lagerheimii*, with all contigs (top) and only the mating-type chromosomes (bottom). The *M. lagerheimii* genome represents a proxy for the ancestral state before recombination suppression and with two separate mating-type chromosomes. The HD mating-type chromosome from *M. lagerheimii* is represented in blue and the PR mating-type chromosome in purple (splitted into two scaffolds), with links to orthologous genes to the *M. superbum* mating-type chromosomes. The positions of the HD and PR mating-type genes are indicated in red and by red links between the two genomes. The PR chromosome shows a substantial increase in size and a chaos of rearrangements.

**Figure S4:**
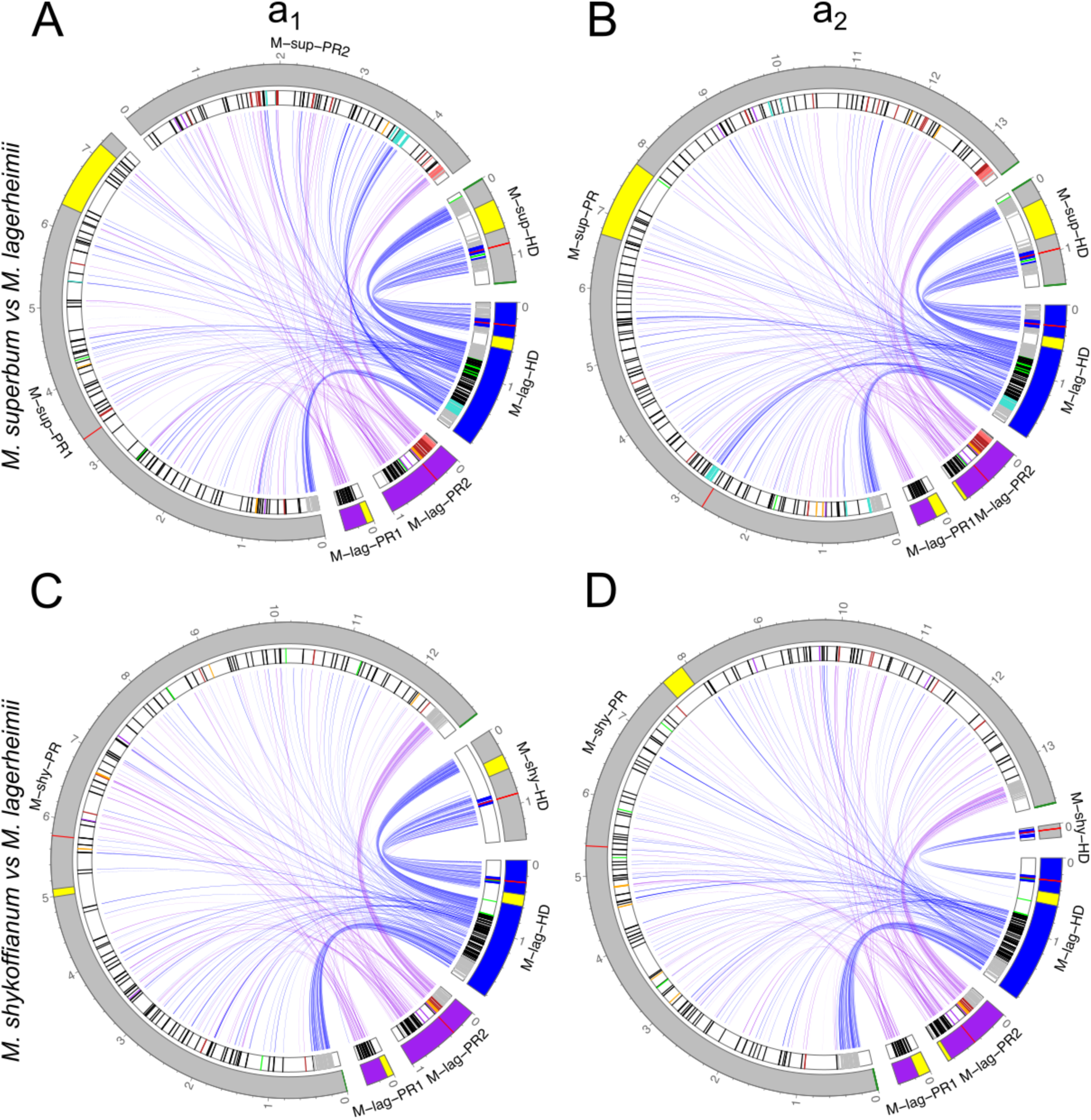
Synteny and rearrangements between the a_1_ (left) and a_2_ (right) HD and PR mating-type chromosomes of *Microbotryum superbum* (A-B) and *Microbotryum shykoffianum* (C-D) compared to those of *M. lagerheimii*. Links are represented between orthologous genes, in blue for the genes on the *M. lagerheimii* HD mating-type chromosome and in purple for the *M. lagerheimii* PR mating-type chromosome. The HD and PR mating-type genes are indicated in red. Centromeres are figured in yellow and telomeres in dark green on the outer track. The strata are indicated by their color on the inner track. Genes that have moved from the ancestral short HD chromosome arm to current PR chromosome, or reciprocally, are in green on the inner track.

**Figure S5:**
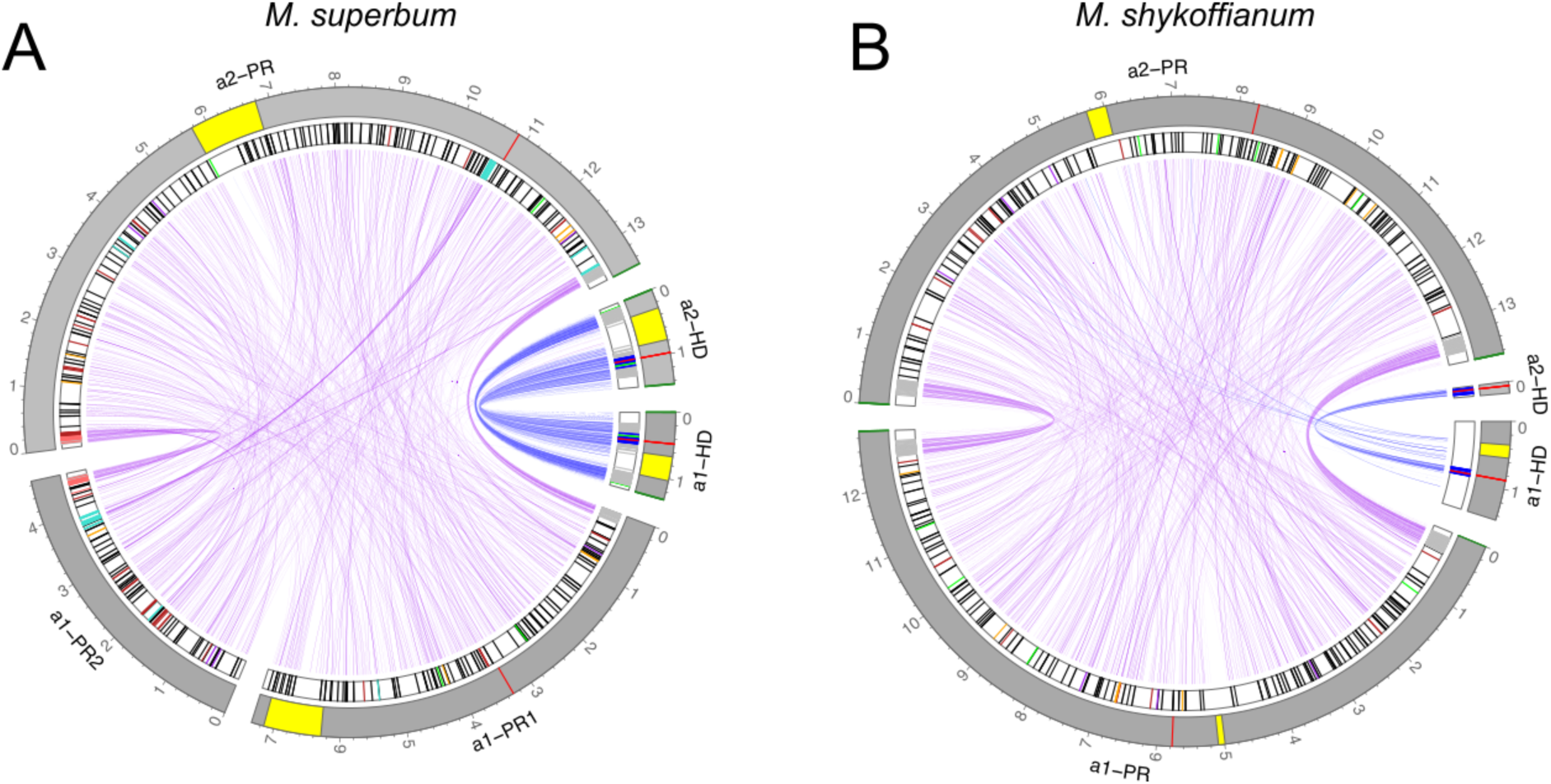
Synteny and rearrangements between the a_1_ and a_2_ mating-type chromosomes (HD and PR chromosomes) of *Microbotryum superbum* (A) and *Microbotryum shykoffianum* (B). Links are represented between orthologous genes, colors corresponding to the evolutionary strata. The HD and PR mating-type genes are indicated in red. Centromeres are figured in yellow and telomeres in dark green on the outer track. The strata are indicated by their color on the inner track. Genes that have moved from ancestral short HD chromosome arm to current PR chromosome, or reciprocally, are in green on the inner track.

**Figure S6:**
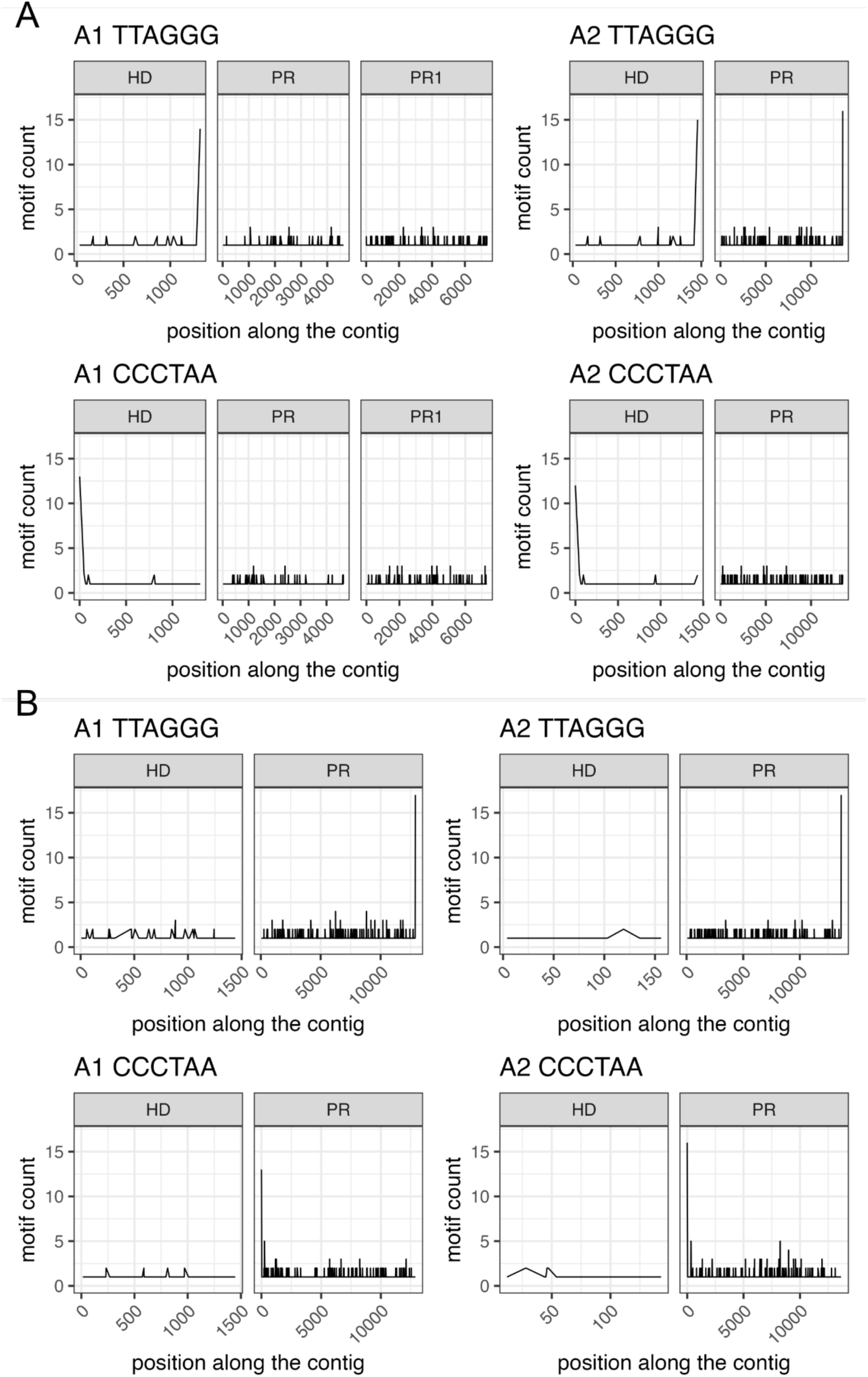
Telomere motifs count over 1kb windows for *Microbotryum violaceum superbum* (A) and *M. shykoffianum* (B). Top panel corresponds to the motif TTAGGG for a_1_ and a_2_ contigs, and bottom panel to the inverse complement CCCTAA for a_1_ and a_2_ contigs.

**Figure S7:**
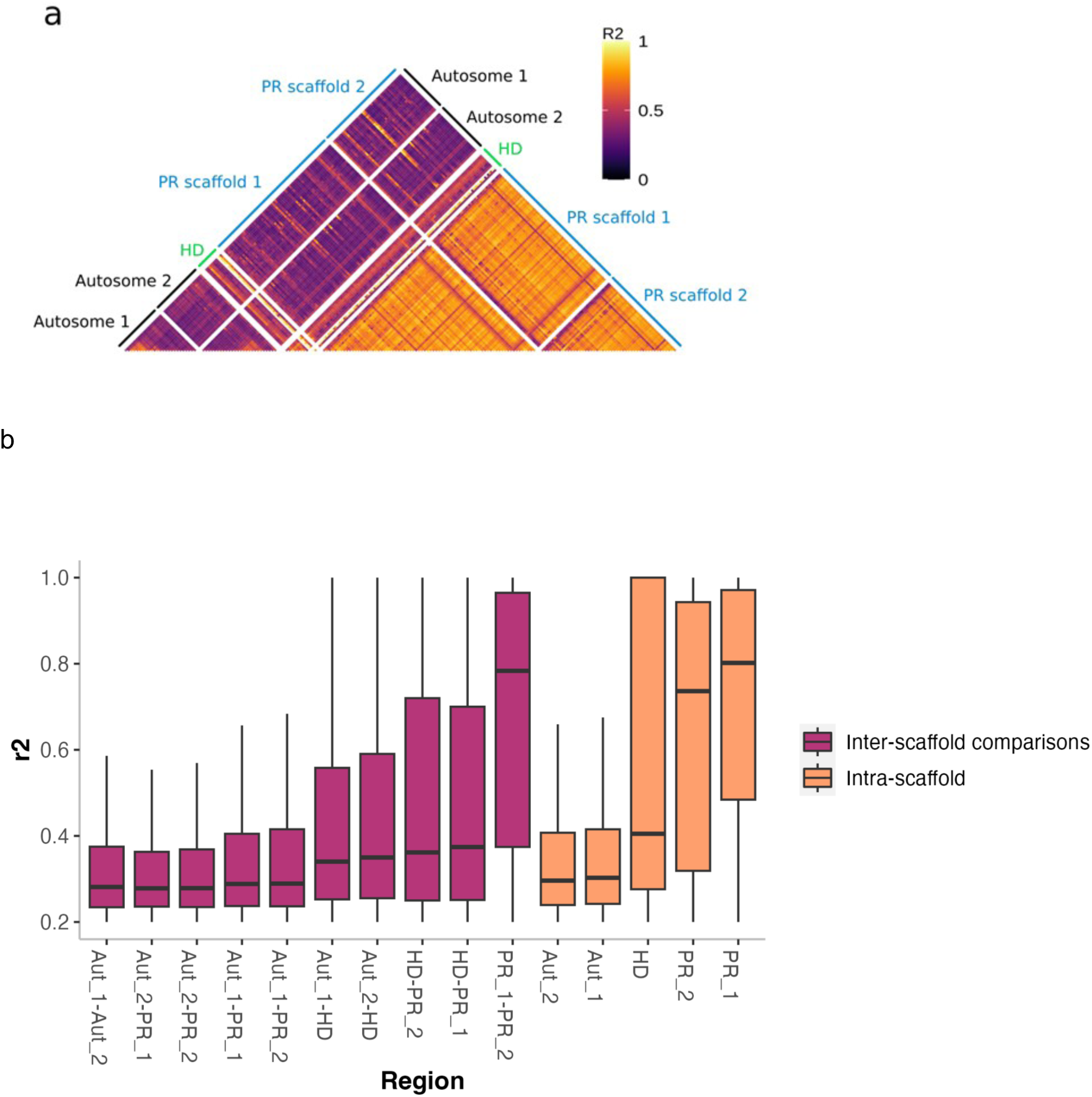
Linkage disequilibrium between the mating-type chromosomes and two autosomes in the a_1_ genome of *Microbotryum superbum.* Linkage disequilibrium as measured by r² is represented with colors along chromosomes (a) and summarized with boxplots (b). The two autosomes are called Aut1 and Aut2, and the two contigs of the PR chromosomes in the a_1_ genome are called PR1 and PR2.

**Figure S8:**
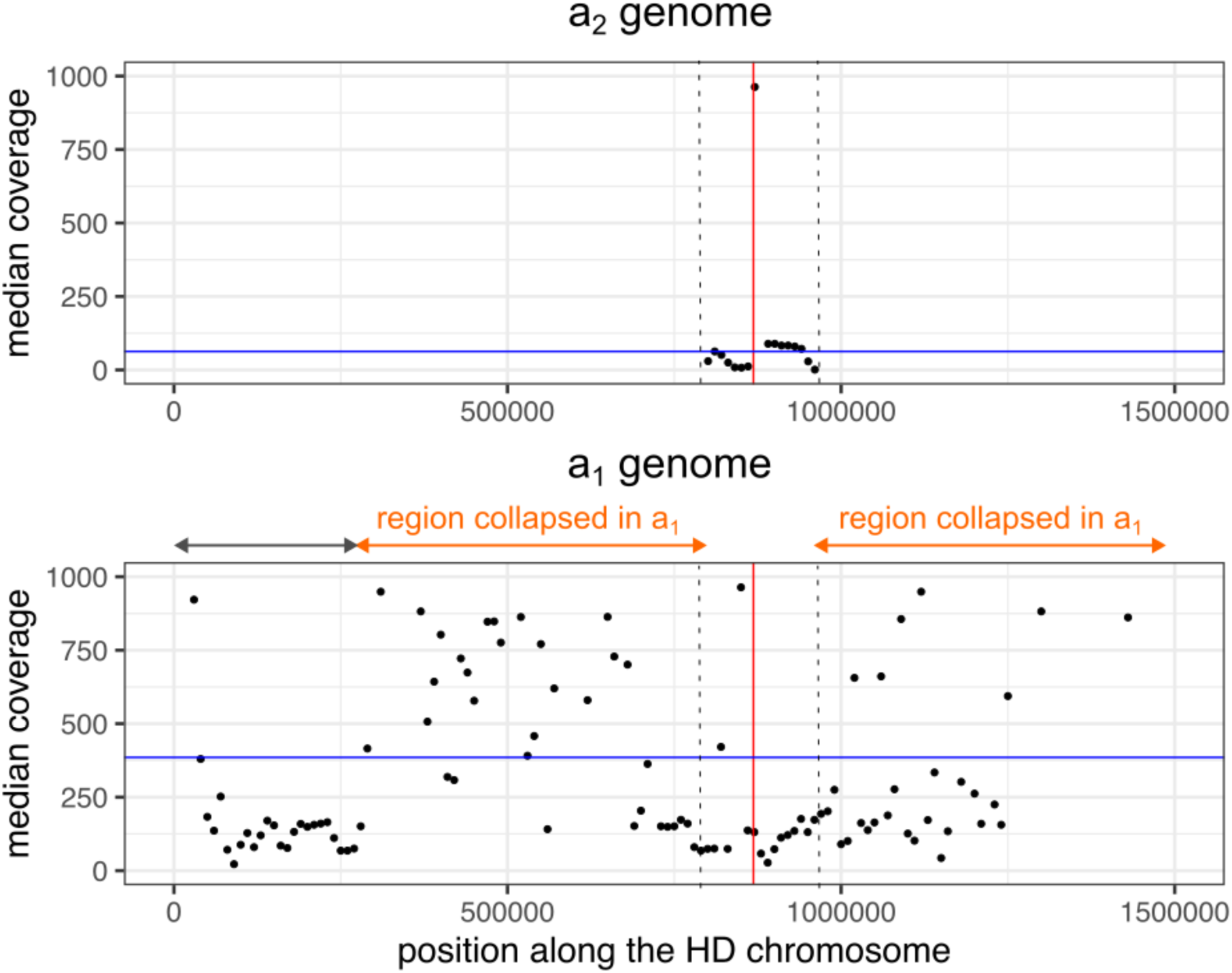
Read mapping coverage along the HD chromosome in *Microbotryum shykoffianum.* The mapping coverage is represented against the a_2_ genome at the top and against the a_1_ genome at the bottom. The location of the canonical HD mating-type genes is represented by the red vertical bar. The median read coverage is indicated by the blue horizontal bar. The orange arrows represent the regions collapsed in the a_1_ genome assembly. The region delimited by vertical dotted lines is the region that was assigned correctly to a_1_ and a_2_ mating types. The grey arrow represents a region that might be missing from the a_2_ genome.

**Figure S9:**
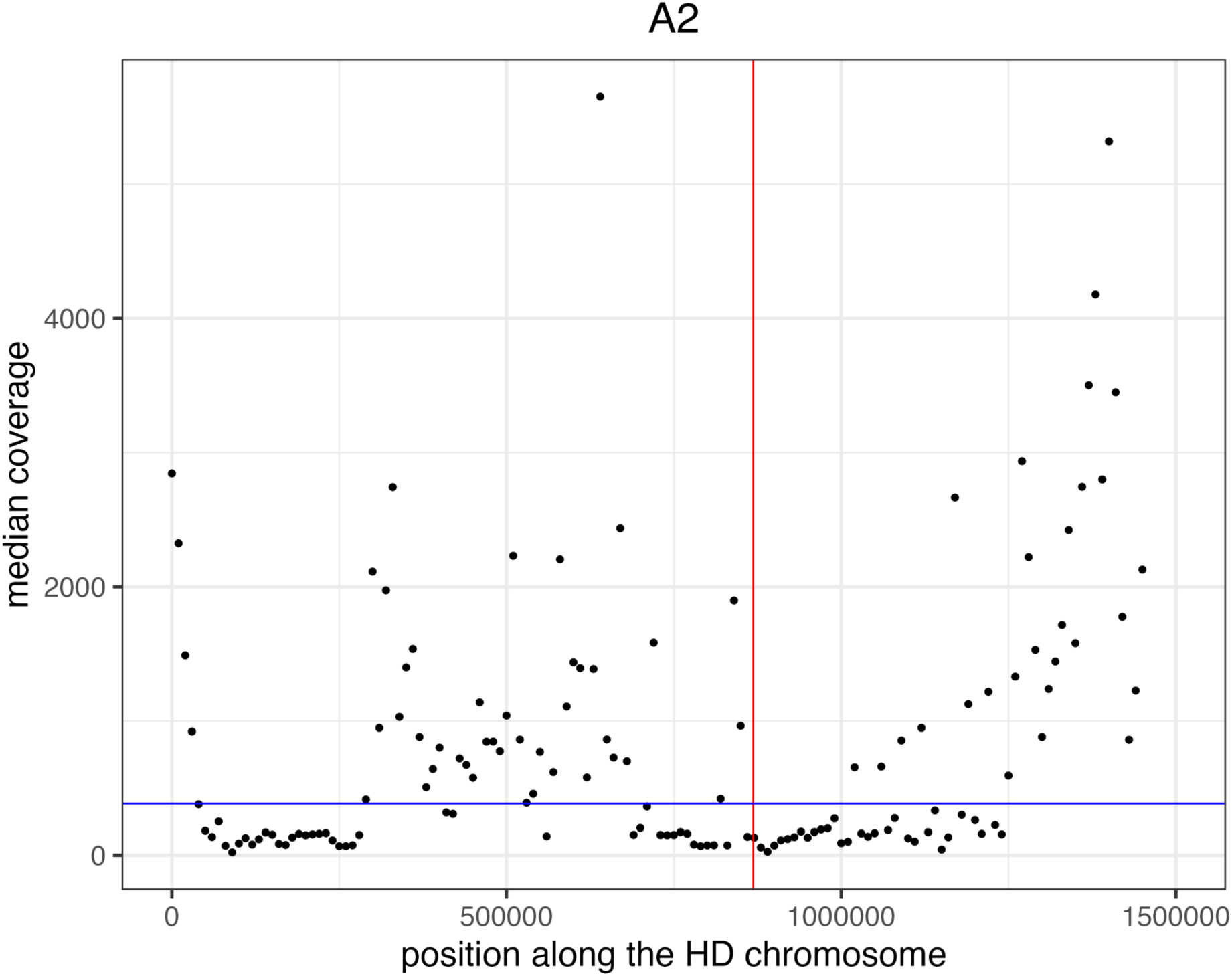
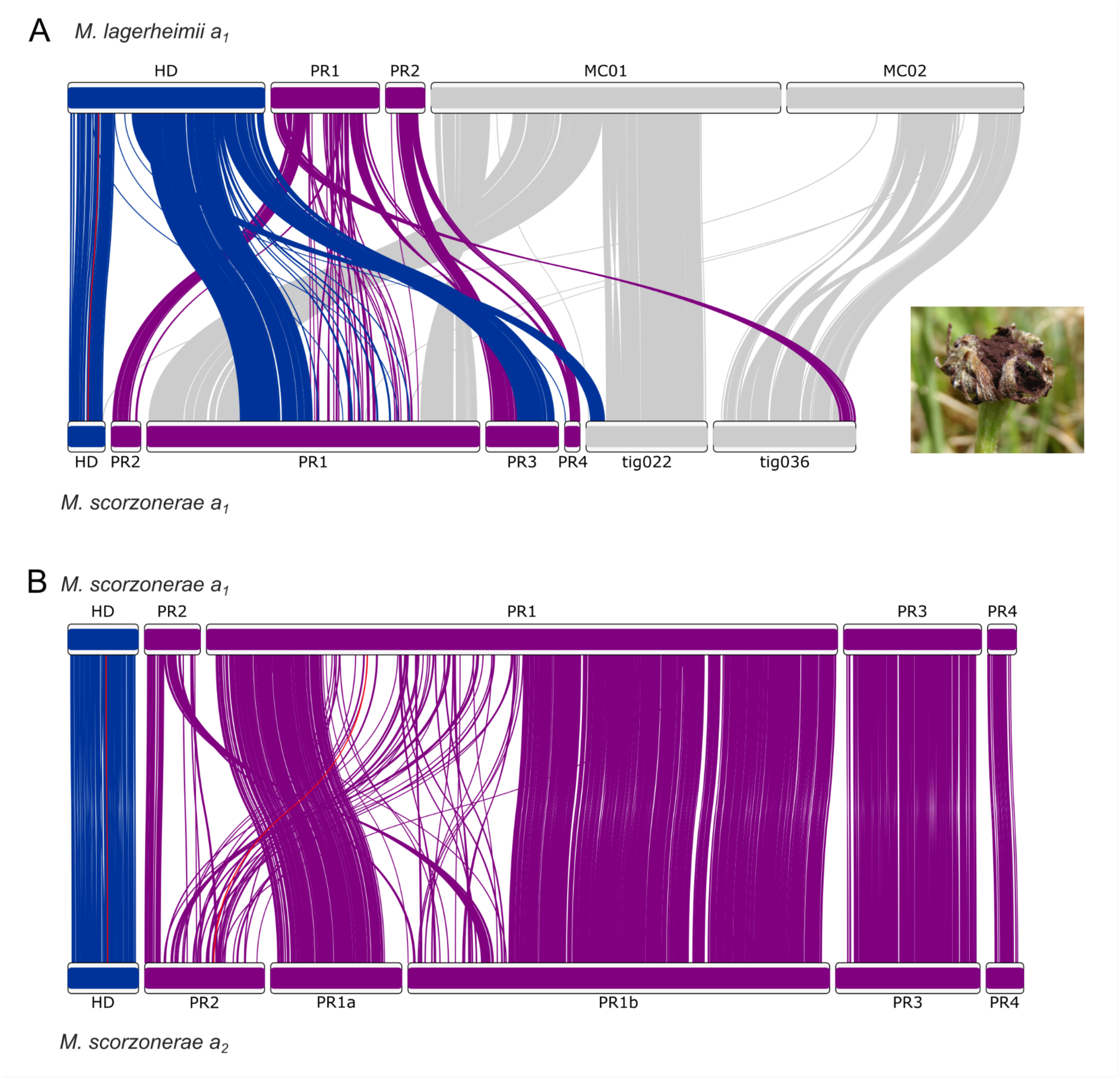
Synteny and rearrangements between the a_1_ genomes of *Microbotryum scorzonarae* and *M. lagerheimii* (A) and between a_1_ and a_2_ genomes of *M. scorzonarae* (B). The *M. lagerheimii* genome represents a proxy for the ancestral state. The HD and PR loci are linked between genomes by red links. On the right, a picture of a diseased flower.

**Figure S10:**
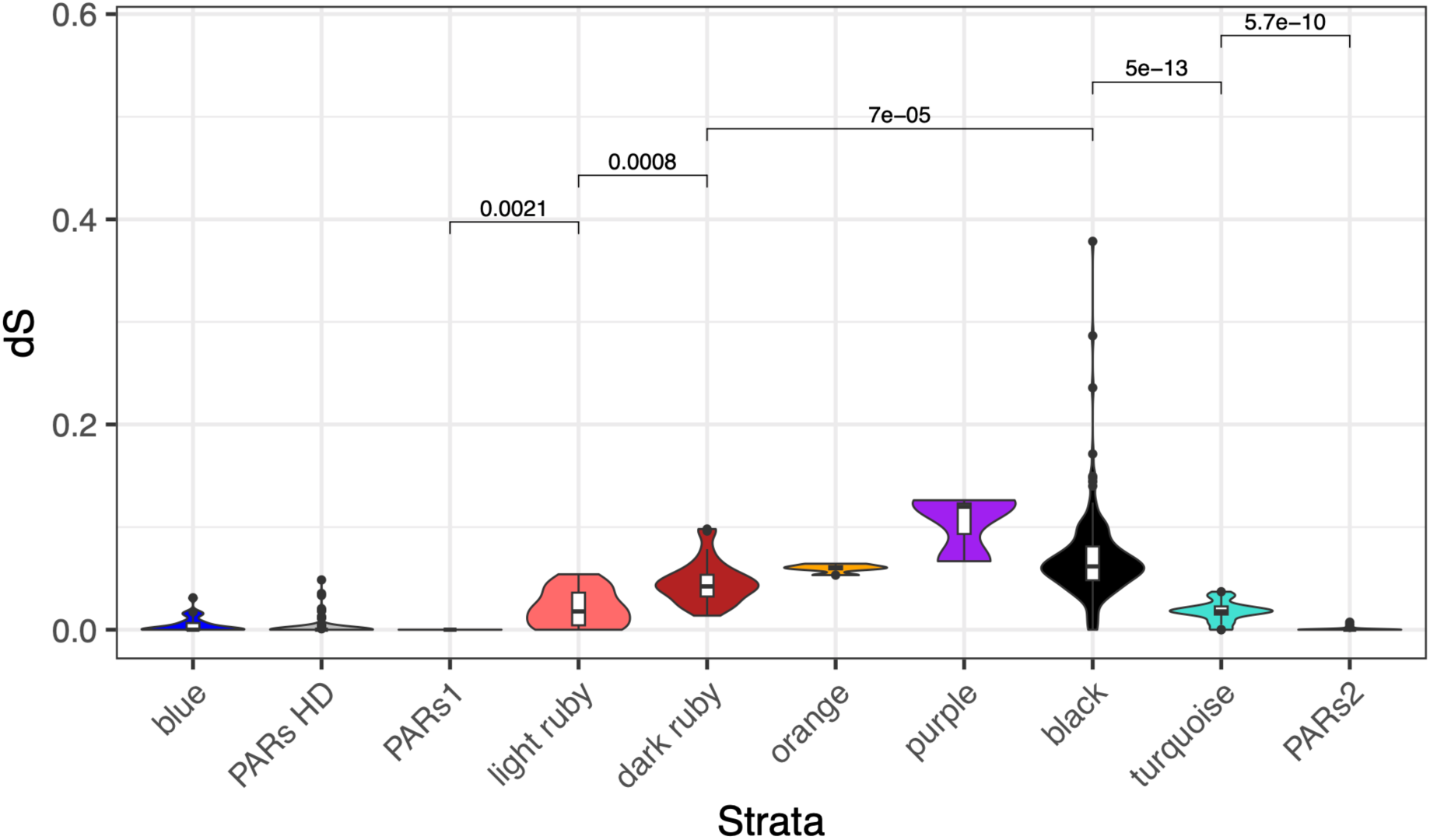
Boxplot of the synonymous divergence (d_S_) values for each stratum in *Microbotryum superbum*. Wilcoxon tests were performed between each pairwise comparison of strata to assess significant differences in mean d_S_ between strata. Only significant p-values are shown.

**Figure S11:**
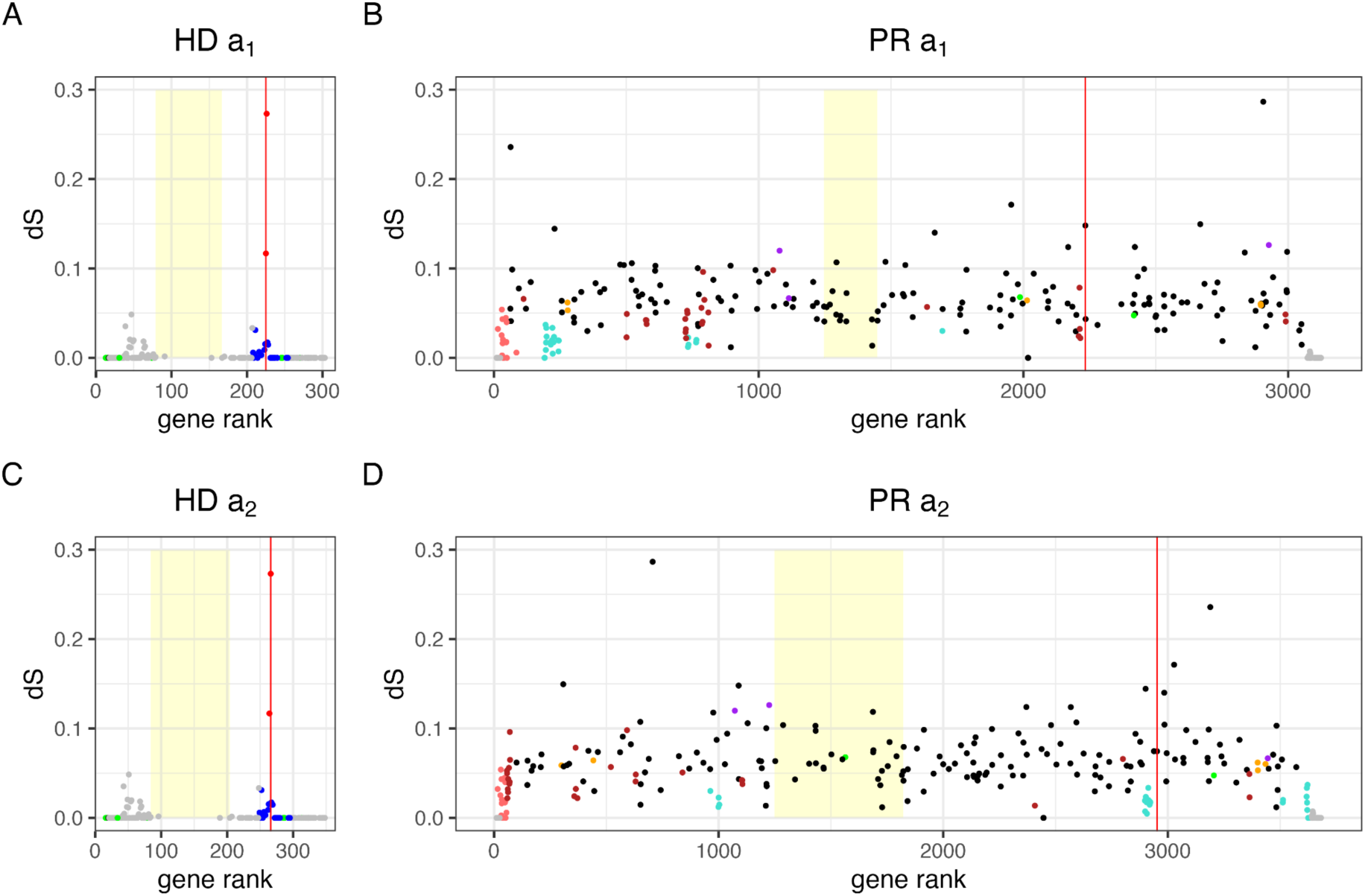
Differentiation and rearrangements in the HD (A-C) and PR (B-D) mating-type chromosomes in *Microbotryum superbum*. A-B correspond to the a_1_ mating type, and C-D to the a_2_ mating type. Per-gene synonymous divergence (d_S_) plotted along the current gene order in the PR and HD chromosomes, in the a_1_ and a_2_ genomes; the points are colored according to their evolutionary stratum assignment: turquoise, light ruby for the *M. superbum* specific evolutionary strata, dark ruby and black for the evolutionary stratum shared by *M. superbum* and *M. shykoffianum,* blue, orange and purple for the ancient evolutionary strata shared by most *Microbotryum* species. The pseudo-autosomal regions (PARs) are in grey. The genes ancestrally located in the alternative mating-type chromosome are in green (i.e. the genes ancestrally in the HD chromosome arm are in green on the PR chromosome, and reciprocally). The yellow boxes correspond to the position of the centromeres. The vertical red lines indicate the position of the HD and PR loci.

**Figure S12:**
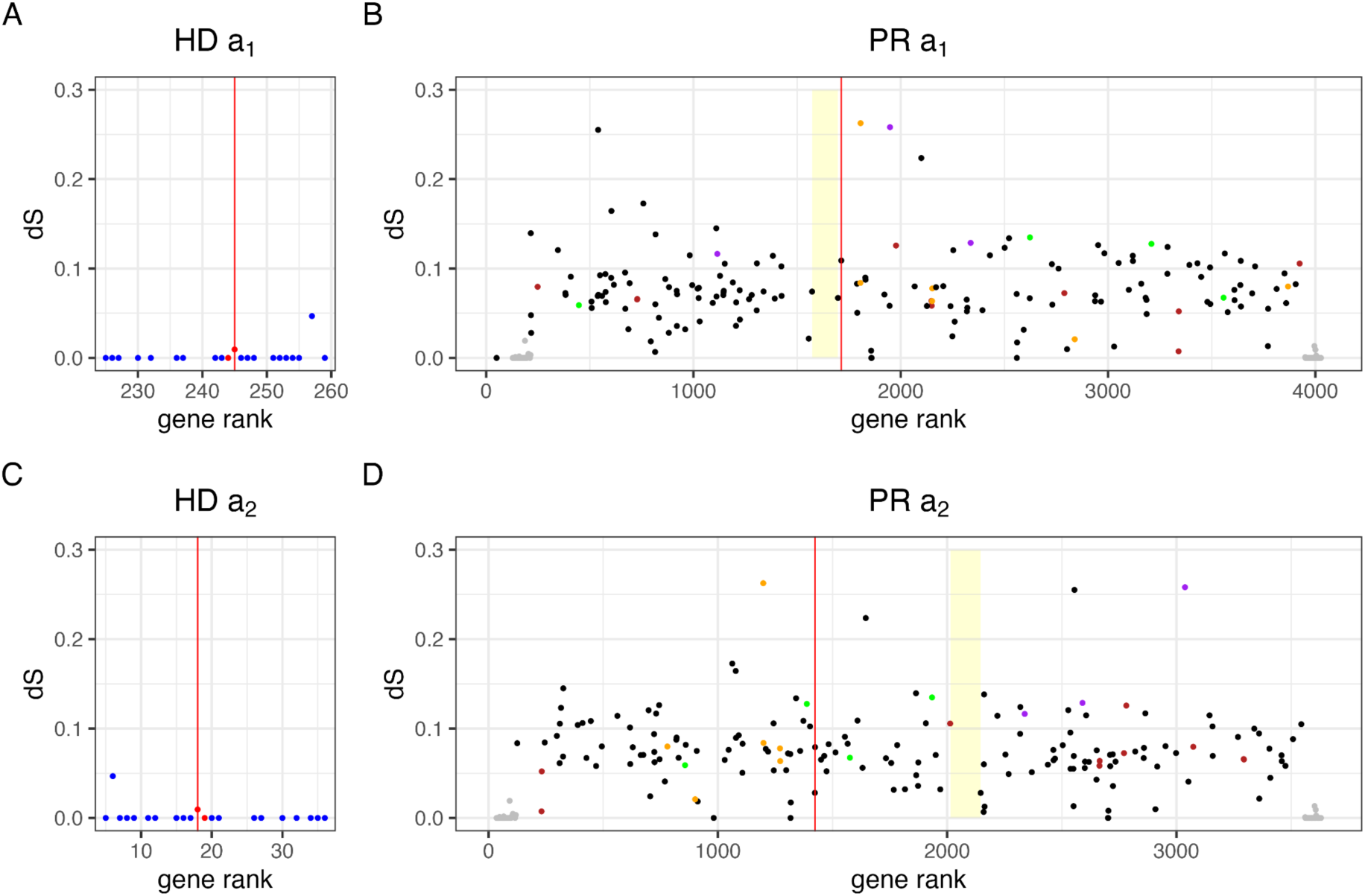
Differentiation and rearrangements in the HD (A-C) and PR (B-D) mating-type chromosomes in *Microbotryum shykoffianum*. A-B correspond to the a_1_ mating type, and C-D to the a_2_ mating type. Per-gene synonymous divergence (d_S_) plotted along the current gene order in the PR and HD chromosomes, in the a_1_ and a_2_ genomes; the points are colored according to their evolutionary stratum assignment: dark ruby and black for the evolutionary strata shared by *M. superbum* and *M. shykoffianum,* blue, orange and purple for the ancient evolutionary strata shared by most *Microbotryum* species. The pseudo-autosomal regions (PARs) are in grey. The genes ancestrally located in the alternative mating-type chromosome are in green (i.e. the genes ancestrally in the HD chromosome arm are in green on the PR chromosome, and reciprocally).

**Figure S13:**
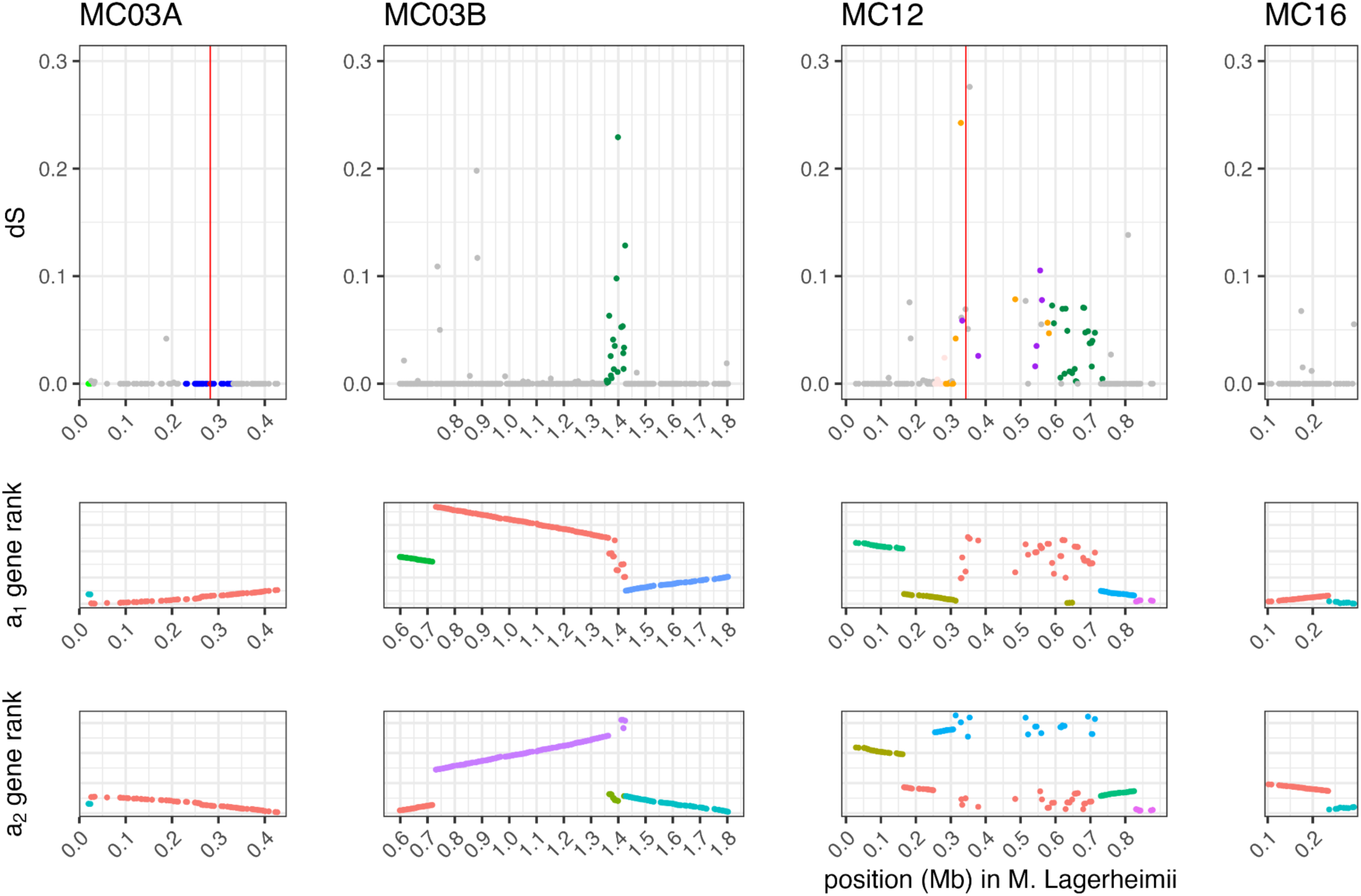
Differentiation and rearrangements between mating-type chromosomes in *Microbotryum scorzoraneae*. **A)** Per-gene synonymous divergence (d_S_) plotted along the ancestral gene order (taking as proxy the gene order along the *M. lagerheimii* mating-type chromosomes, its HD chromosomes encompassing the MC03A and B contigs and the PR chromosome the MC12 and MC16 contigs). The points are colored according to their evolutionary stratum assignment: emerald and quartz for the young evolutionary stratum specific to *M. scorzonareae,* and blue, orange and purple for the ancient evolutionary strata shared by most *Microbotryum* species. The pseudo-autosomal regions (PARs) are in grey. **B)** Rearrangements compared to the ancestral gene order figured by plotting the gene rank in the current gene order (in the a_1_ genome and in the a_2_ genome) as a function of the gene rank in the ancestral gene order. The different contigs in *M. scorzonareae* have different colors in the gene rank panels.

**Figure S14-.**
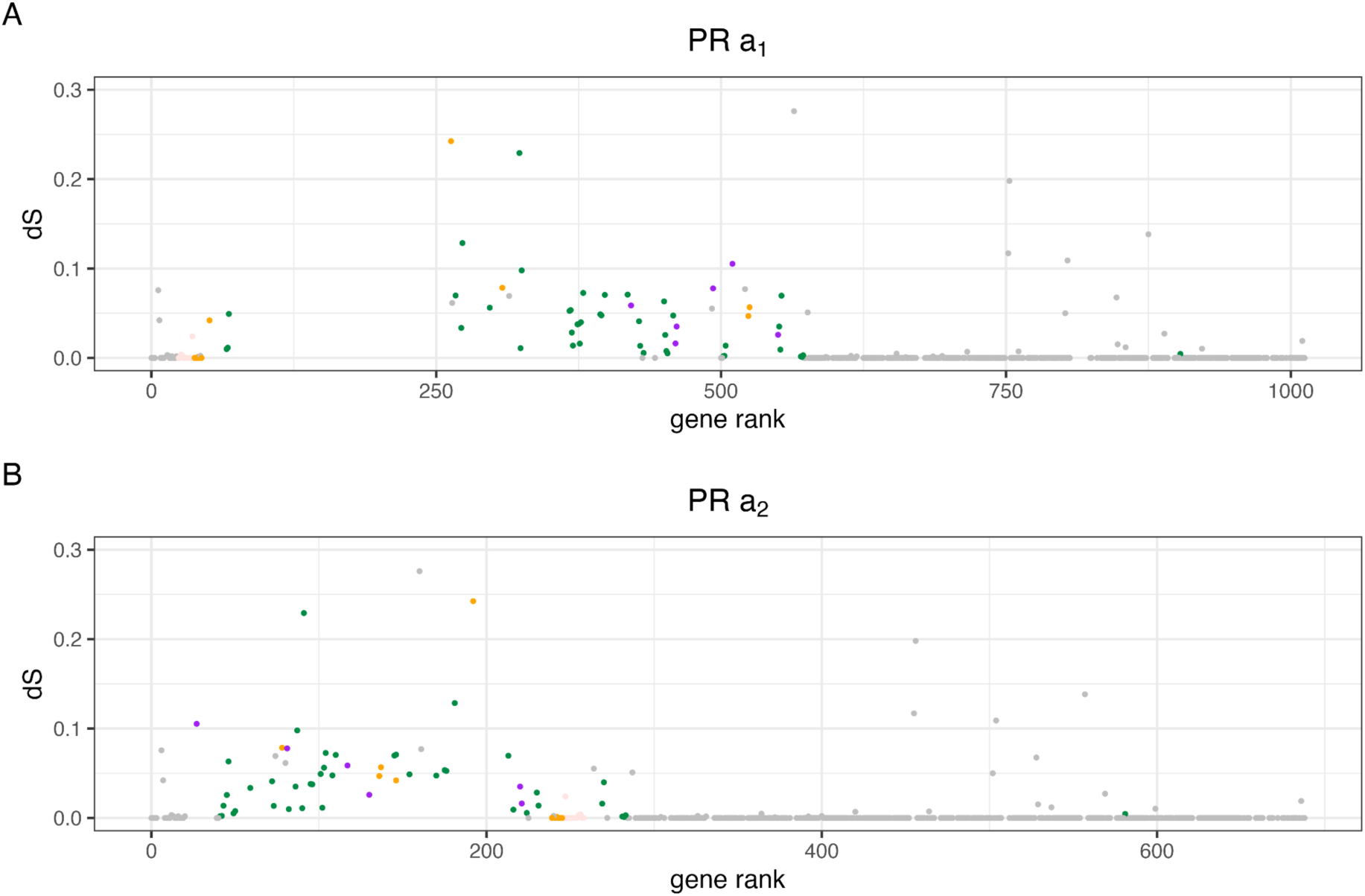
Differentiation and rearrangements in the PR mating-type chromosomes in *Microbotryum scorzoraneae*. Per-gene synonymous divergence (d_S_) plotted along the current gene order in the PR chromosome in the a_1_ (A) and a_2_ (B) genomes; the points are colored according to their evolutionary stratum assignment: emerald and quartz rose or the *M. scorzoraneae* specific evolutionary strata, orange and purple for the ancient evolutionary strata shared by most *Microbotryum* species. The pseudo-autosomal regions (PARs) are in grey.

## Supplementary Tables

**Table S1:**
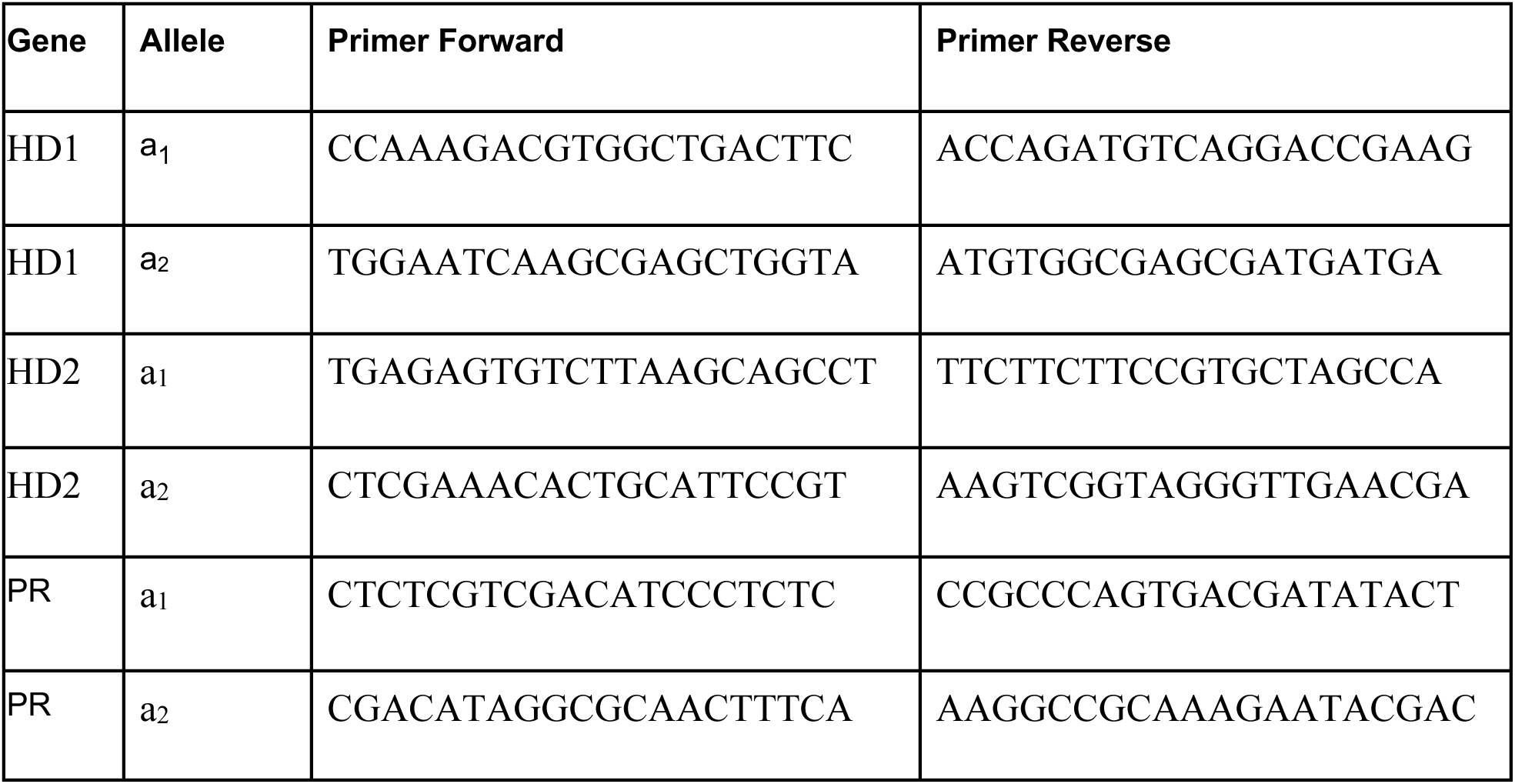
Primer pairs designed for specifically amplifying the different alleles at the genes HD1, HD2 and PR as designed from *Microbotryum superbum* 1065.

**Table S2:** Segregation of mating-type alleles in six tetrads of *Microbotryum superbum* (two opposite cells in a linear tetrad were analysed, i.e. P and D, corresponding to segregation at first meiotic division). A + sign indicates amplification by PCR of a fragment using primers designed to amplify specifically the a_1_ or a_2_ alleles in the *M. superbum* reference genome for the three mating-type genes PR, HD1 and HD2.

| Tetrad | Cell in tetrad | PR a<sub>1</sub> | PR a<sub>2</sub> | HD1 a<sub>1</sub> | HD1 a<sub>2</sub> | HD2 a<sub>1</sub> | HD2 a<sub>2</sub> |
| --- | --- | --- | --- | --- | --- | --- | --- |
| 12 | P | + | - | - | + | - | + |
| 12 | D | - | + | - | + | - | + |
| 14 | P | - | + | + | - | + | - |
| 14 | D | + | - | - | + | - | + |
| 6 | P | - | + | - | + | - | - |
| 6 | D | + | - | - | + | - | - |
| 10 | P | + | - | + | - | + | - |
| 10 | D | - | + | + | - | + | - |
| 6 | P | - | + | - | + | - | + |
| 6 | D | + | - | - | + | - | + |
| 3 | P | + | - | + | - | + | - |
| 3 | D | - | + | - | + | - | + |

**Table S3:** RNA seq sources (see excel file)

## Notes

### Competing Interest Statement

The authors have declared no competing interest.

### Summary of Updates

Minor revisions on material and methods and results. Authors list updated.

## REFERENCES

1 Lande, R. & Schemske, D. The evolution of self-fertilization and inbreeding depression in plants. I. Genetic models. Evolution 39, 24–40 (1985).

2 Charlesworth, D. & Charlesworth, B. Inbreeding depression and its evolutionary consequences. Annu Rev Ecol Evol Syst 18, 237–268 (1987).

3 Charlesworth, D., Morgan, M. & Charlesworth, B. Inbreeding depression, genetic load, and the evolution of outcrossing rates in a multilocus system with no linkage. Evolution 44, 1469–1489 (1990).

4 Charlesworth, D. Plant sex determination and sex chromosomes. Heredity 88, 94–101 (2002).

5 Igic, B., Lande, R. & Kohn, J. Loss of self-incompatibility and its evolutionary consequences. Int J Plant Sci 169, 93–104 (2008).

6 Hereford, J. Does selfing or outcrossing promote local adaptation?. Am J Bot 97, 298–302 (2010).

7 Lande, R. Evolution of phenotypic plasticity in colonizing species. Mol Ecol 24, 2038–2045 (2015).

8 Nieuwenhuis, B. P. S. et al. Evolution of uni- and bifactorial sexual compatibility systems in fungi. Heredity 111, 445–455, doi:10.1038/hdy.2013.67 (2013).

9 Goldberg, E. & Igić, B. Tempo and mode in plant breeding system evolution. Evolution 66, 3701–3709 (2012).

10 Chantha, S., Herman, A., Platts, A., Vekemans, X. & Schoen, D. Secondary evolution of a self-incompatibility locus in the Brassicaceae genus *Leavenworthia*. PLoS Biol 11, e1001560 (2013).

11 Hanschen, E., Herron, M., Wiens, J., Nozaki, H. & Michod, R. Repeated evolution and reversibility of self-fertilization in the volvocine green algae. Evolution 72, 386–398 (2018).

12 Beukeboom, L. & Perrin, N. 92-93 (Oxford University Press, Oxford, 2014).

13 Vekemans, X., Poux, C., Goubet, P. & Castric, V. The evolution of selfing from outcrossing ancestors in Brassicaceae: What have we learned from variation at the S-locus?. J Evol Biol 27, 1372–1385 (2014).

14 Billiard, S. et al. Having sex, yes, but with whom? Inferences from fungi on the evolution of anisogamy and mating types. Biol Rev 86, 421–442 (2011).

15 Billiard, S., López-Villavicencio, M., Hood, M. & Giraud, T. Sex, outcrossing and mating types: unsolved questions in fungi and beyond. J Evol Biol 25, 1020–1038 (2012).

16 Coelho, M. A., Bakkeren, G., Sun, S., Hood, M. E. & Giraud, T. Fungal Sex: The Basidiomycota. Microbiology Spectrum 5, 1–30, doi:10.1128/microbiolspec.FUNK-0046-2016 (2017).

17 Branco, S. et al. Evolutionary strata on young mating-type chromosomes despite the lack of sexual antagonism. Proceedings of the National Academy of Sciences 114, 7067–7072, doi:10.1073/pnas.1701658114 (2017).

18 Branco, S. et al. Multiple convergent supergene evolution events in mating-type chromosomes. Nature Communications 9, 2000, doi:10.1038/s41467-018-04380-9 (2018).

19 Charlesworth, D. The status of supergenes in the 21st century: recombination suppression in Batesian mimicry and sex chromosomes and other complex adaptations. Evolutionary Applications 9, 74–90, doi:10.1111/eva.12291 (2016).

20 Charlesworth, D. Evolution of recombination rates between sex chromosomes. Philosophical Transactions of the Royal Society B-Biological Sciences 372, doi:10.1098/rstb.2016.0456 (2017).

21 Charlesworth, D. When and how do sex-linked regions become sex chromosomes? Evolution 75, 569–581, doi:10.1111/evo.14196 (2021).

22 Bergero, R. & Charlesworth, D. The evolution of restricted recombination in sex chromosomes. Trends in Ecology & Evolution 24, 94–102, doi:10.1016/j.tree.2008.09.010 (2009).

23 Hartmann, F. E. et al. Recombination suppression and evolutionary strata around mating-type loci in fungi: documenting patterns and understanding evolutionary and mechanistic causes. New Phytologist 229, 2470–2491, doi:10.1111/nph.17039 (2021).

24 Vittorelli, N. et al. Stepwise recombination suppression around the mating-type locus in the fungus *Schizothecium tetrasporum* (Ascomycota, Sordariales). PLoS Genetics 19, e1010347 (2022).

25 Ironside, J. E. No amicable divorce? Challenging the notion that sexual antagonism drives sex chromosome evolution. Bioessays 32, 718–726, doi:10.1002/bies.200900124 (2010).

26 Charlesworth, D., Charlesworth, B. & Marais, G. Steps in the evolution of heteromorphic sex chromosomes. Heredity 95 118–128 (2005).

27 Wright, A. E., Dean, R., Zimmer, F. & Mank, J. E. How to make a sex chromosome. Nature Communications 7, 12087, doi:10.1038/ncomms12087 (2016).

28 Bazzicalupo, A. L., Carpentier, F., Otto, S. P. & Giraud, T. Little evidence of antagonistic selection in the evolutionary strata of fungal mating-type chromosomes *(Microbotryum lychnidis-dioicae)*. G3-Genes Genomes Genetics 9, 1987-1998, doi:10.1534/g3.119.400242 (2019).

29 Jeffries, D. L., Gerchen, J. F., Scharmann, M. & Pannell, J. R. A neutral model for the loss of recombination on sex chromosomes. Philosophical Transactions of the Royal Society B-Biological Sciences 376, doi:10.1098/rstb.2020.0096 (2021).

30 Kent, T. V., Uzunovic, J. & Wright, S. I. Coevolution between transposable elements and recombination. Philosophical Transactions of the Royal Society B-Biological Sciences 372, doi:10.1098/rstb.2016.0458 (2017).

31 Jay, P., Tezenas, E., Véber, A. & Giraud, T. Sheltering of deleterious mutations explains the stepwise extension of recombination suppression on sex chromosomes and other supergenes. PLoS Biology 20, e3001698, doi:10.1371/journal.pbio.3001698 (2022).

32 Lenormand, T. & Roze, D. Y recombination arrest and degeneration in the absence of sexual dimorphism. Science 375, 663-+, doi:10.1126/science.abj1813 (2022).

33 Jay, P., Jeffries, D., Hartmann, F. E., Véber, A. & Giraud, T. Why do sex chromosomes progressively lose recombination?. Trends in Genetics in press (2024).

34 Hill, W. G. & Robertson, A. The effect of linkage on limits to artificial selection. Genetics Research 8, 269–294, doi:10.1017/S001667230800949X (1966).

35 Muller, H. J. Some genetic aspects of sex. The American Naturalist 66, 118–138 (1932).

36 Charlesworth, B. & Charlesworth, D. The degeneration of Y chromosomes. Philosophical Transactions of the Royal Society B-Biological Sciences 355, 1563–1572, doi:10.1098/rstb.2000.0717 (2000).

37 Charlesworth, D. The timing of genetic degeneration of sex chromosomes. Phil. Trans. R. Soc. B 376, 20200093 (2021).

38 Bachtrog, D. Y-chromosome evolution: emerging insights into processes of Y-chromosome degeneration. Nature Reviews Genetics 14, 113–124, doi:10.1038/nrg3366 (2013).

39 Carpentier, F. et al. Tempo of degeneration across independently evolved nonrecombining regions. Molecular Biology and Evolution 39, doi:10.1093/molbev/msac060 (2022).

40 Fontanillas, E. et al. Degeneration of the nonrecombining regions in the mating-type chromosomes of the anther-smut fungi. Molecular Biology and Evolution 32, 928–943, doi:10.1093/molbev/msu396 (2015).

41 Duhamel, M. et al. Onset and stepwise extensions of recombination suppression are common in mating-type chromosomes of *Microbotryum* anther-smut fungi. Journal of Evolutionary Biology 35, 1619–1634, doi:10.1111/jeb.13991 (2022).

42 Badouin, H. et al. Chaos of rearrangements in the mating-type chromosomes of the anther-smut fungus *Microbotryum lychnidis-dioicae*. Genetics 200, 1275–1284, doi:10.1534/genetics.115.177709 (2015).

43 43 Kües, U., James, T. Y. & Heitman, J. in Evolution of Fungi and Fungal-Like Organisms (eds S. Pöggeler & J. Wöstemeyer) Pp. 97–160 (Springer, 2011).

44 Bakkeren, G. & Kronstad, J. W. Linkage of mating-type loci distinguishes bipolar from tetrapolar mating in basidiomycetous smut fungi. Proceedings of the National Academy of Sciences of the United States of America 91, 7085–7089, doi:10.1073/pnas.91.15.7085 (1994).

45 Xu, J. et al. Dandruff-associated *Malassezia* genomes reveal convergent and divergent virulence traits shared with plant and human fungal pathogens. Proceedings of the National Academy of Sciences of the United States of America 104, 18730–18735, doi:10.1073/pnas.0706756104 (2007).

46 Coelho, M. A., Ianiri, G., David-Palmaa, M. & Heitman, J. Frequent transitions in mating-type locus chromosomal organization in Malassezia and early steps in sexual reproduction. PNAS 120, e2305094120 (2023).

47 James, T. Y., Srivilai, P., Kues, U. & Vilgalys, R. Evolution of the bipolar mating system of the mushroom *Coprinellus disseminatus* from its tetrapolar ancestors involves loss of mating-type-specific pheromone receptor function. Genetics 172, 1877–1891, doi:10.1534/genetics.105.051128 (2006).

48 Chen, B. Z. et al. Fruiting body formation in *Volvariella volvacea* can occur independently of its MAT-a-controlled bipolar mating system, enabling homothallic and heterothallic life cycles. G3-Genes Genomes Genetics 6, 2135-2146, doi:10.1534/g3.116.030700 (2016).

49 Kues, U. et al. A chimeric homeodomain protein causes self-incompatibility and constitutive sexual development in the mushroom *Coprinus cinereus*. Embo Journal 13, 4054–4059, doi:10.1002/j.1460-2075.1994.tb06722.x (1994).

50 Hood, M. E. et al. Distribution of the anther-smut pathogen non species of the Caryophyllaceae. New Phytologist 187, 217–229, doi:10.1111/j.1469-8137.2010.03268.x (2010).

51 Hartmann, F. E. et al. in Annual Review of Phytopathology, Vol 57, 2019 Vol. 57 Annual Review of Phytopathology (eds J. E. Leach & S. E. Lindow) 431-457 (2019).

52 Le Gac, M., Hood, M. E., Fournier, E. & Giraud, T. Phylogenetic evidence of host-specific cryptic species in the anther smut fungus. Evolution 61, 15–26, doi:10.1111/j.1558-5646.2007.00002.x (2007).

53 Refrégier, G. et al. Cophylogeny of the anther smut fungi and their caryophyllaceous hosts: Prevalence of host shifts and importance of delimiting parasite species for inferring cospeciation. BMC Evolutionary Biology 8, doi:10.1186/1471-2148-8-100 (2008).

54 Hood, M. E., Antonovics, J. & Koskella, B. Shared forces of sex chromosome evolution in haploids and diploids. Genetics 168, 141–146 (2004).

55 Feurtey, A. et al. Strong phylogeographic co-structure between the anther-smut fungus and its white campion host. New Phytologist 212, 668–679, 10.1111/nph.14125 (2016).

56 Giraud, T., Yockteng, R., Lopez-Villavicencio, M., Refregier, G. & Hood, M. E. The mating system of the anther smut fungus, Microbotryum violaceum: selfing under heterothallism. Euk. Cell 7, 765–775 (2008).

57 Hartmann, F. E. et al. Congruent population genetic structures and divergence histories in anther-smut fungi and their host plants Silene italica and the Silene nutans species complex. Molecular Ecology 29, 1154–1172, 10.1111/mec.15387 (2020).

58 Hood, M. E. & Antonovics, J. Intratetrad mating, heterozygosity, and the maintenance of deleterious alleles in *Microbotryum violaceum (= Ustilago violacea)*. Heredity 85, 231–241, doi:10.1046/j.1365-2540.2000.00748.x (2000).

59 Alexander, H. M., Thrall, P. H., Antonovics, J., Jarosz, A. M. & Oudemans, P. V. Population dynamics and genetics of plant disease: a case study of anther-smut disease. Ecology 77, 990–996 (1996).

60 Antonovics, J., Iwasa, Y. & Hassel, M. P. A generalized model of parasitoid, veneral, and vector-based transmission processes. Am. Nat. 145, 661–675 (1995).

61 Hood, M. E., Antonovics, J. & Heishman, H. Karyotypic similarity identifies multiple host-shifts of a pathogenic fungus in natural populations. Infection, Genetics and Evolution 2, 167–172 (2003).

62 Hartmann, F. E. et al. Higher gene flow in sex-related chromosomes than in autosomes during fungal divergence. Molecular Biology and Evolution 37, 668–682, doi:10.1093/molbev/msz252 (2020).

63 Zakharov, I. A. [Intratetrad mating and its genetic and evolutionary consequences]. Genetika 41, 508–519 (2005).

64 Zakharov, I. A. Intratetrad mating as the driving force behind the formation of sex chromosomes in fungi. Trends in Genetics and Evolution 6, doi: 10.24294/tge.v24296i24291.24252 (2023).

65 Carpentier, F. et al. Convergent recombination cessation between mating-type genes and centromeres in selfing anther-smut fungi. Genome Research 29, 944–953, doi:10.1101/gr.242578.118.5 (2019).

66 Thomas, A., Shykoff, J., Jonot, O. & Giraud, T. Sex-ratio bias in populations of the phytopathogenic fungus Microbotryum violaceum from several host species. International Journal of Plant Sciences 164, 641–647, doi:10.1086/375374 (2003).

67 Kaltz, O. & Shykoff, J. A. Sporidial mating-type ratios of teliospores from natural populations of the anther smut fungus *Microbotryum* (equals *Ustilago) violaceum*. International Journal of Plant Sciences 158, 575–584, doi:10.1086/297470 (1997).

68 Oudemans, P. V. et al. The distribution of mating-type bias in natural populations of the anther-smut Ustilago violacea on Silene alba in Virginia. Mycologia 90, 372–381, doi:10.2307/3761395 (1998).

69 Duhamel, M., Hood, M. E., Rodriguez de la Vega, R. C. & Giraud, T. Dynamics of transposable element accumulation in the non-recombining regions of mating-type chromosomes in anther-smut fungi. Nature Communications 14, 5692 (2023).

70 Horns, F., Petit, E. & Hood, M. E. Massive expansion of *Gypsy*-Like retrotransposons in *Microbotryum* fungi. Genome Biology and Evolution 9, 363–371, doi:10.1093/gbe/evx011 (2017).

71 Hood, M. E. & Antonovics, J. Mating Within the Meiotic Tetrad and the Maintenance of Genomic. 1759, 1751–1759 (2004).

72 Devier, B., Aguileta, G., Hood, M. & Giraud, T. Ancient trans-specific polymorphism at pheromone receptor genes in basidiomycetes. Genetics 181, 209–223 (2009).

73 Day, A. & Jones, J. The production and characteristics of diploids in *Ustilago violacea*. Genet. Res., Camb. 11, 63–81 (1968).

74 Gladieux, P. et al. Maintenance of fungal pathogen species that are specialized to different hosts: allopatric divergence and introgression through secondary contact. Molecular Biology and Evolution 28, 459–471, doi:10.1093/molbev/msq235 (2011).

75 Lee, N., Bakkeren, G., Wong, K., Sherwood, J. E. & Kronstad, J. W. The mating-type and pathogenicity locus of the fungus Ustilago hordei spans a 500-kb region. Proceedings of the National Academy of Sciences of the United States of America 96, 15026–15031, doi:10.1073/pnas.96.26.15026 (1999).

76 Fraser, J. A. et al. Convergent evolution of chromosomal sex-determining regions in the animal and fungal kingdoms. Plos Biol. 2, 2243–2255 (2004).

77 Rabe, F. et al. A complete toolset for the study of *Ustilago bromivora* and *Brachypodium sp* as a fungal-temperate grass pathosystem. Elife 5, doi:10.7554/eLife.20522 (2016).

78 Sun, S., Coelho, M. A., Heitman, J. & Nowrousian, M. Convergent evolution of linked mating-type loci in basidiomycete fungi. PLoS Genet. 15, e1008365 (2019).

79 Yi, R. et al. Genomic structure of the A mating-type locus in a bipolar basidiomycete, Pholiota nameko. Mycol Res. 113, 240–248 (2009).

80 Wang, Y.-W. et al. Invasive Californian death caps develop mushrooms unisexually and bisexually. Nature Communications 14, 6560 (2023).

81 Cabrita, A. et al. Multiple pathways to homothallism in closely related yeast lineages in the Basidiomycota. mBio 12, 10.1128/mbio.03130-03120, doi:10.1128/mbio.03130-20 (2021).

82 de Vienne, D. M., Hood, M. E. & Giraud, T. Phylogenetic determinants of potential host shifts in fungal pathogens. Journal of Evolutionary Biology 22, 2532–2541, doi:10.1111/j.1420-9101.2009.01878.x (2009).

83 Petit, E. et al. Co-occurrence and hybridization of anther-smut pathogens specialized on Dianthus hosts. Molecular Ecology 26, 1877–1890, 10.1111/mec.14073 (2017).

84 Denchev, C. M., Giraud, T. & Hood, M. E. Three new species of anthericolous smut fungi on Caryophyllaceae. Mycologica Balcanica 6, 79–84 (2009).

85 Liu, T., Tian, H., He, S. & Guo, L. *Microbotryum scorzonerae* (Microbotryaceae), new to China, on a new host plant. Mycotaxon 108, 245–247 (2009).

86 Le Gac, M., Hood, M. E. & Giraud, T. Evolution of reproductive isolation within a parasitic fungal species complex. Evolution 61, 1781–1787, doi:10.1111/j.1558-5646.2007.00144.x (2007).

87 Perlin, M. H. et al. Sex and parasites: Genomic and transcriptomic analysis of *Microbotryum lychnidis-dioicae*, the biotrophic and plant-castrating anther smut fungus. BMC genomics 16, 1–24, doi:10.1186/s12864-015-1660-8 (2015).

88 San Toh, S., et al. Pas de deux: an Intricate dance of anther smut and its host. 8, 505–518, doi:10.1534/g3.117.300318 (2018).

89 Li, H. Minimap2: pairwise alignment for nucleotide sequences. Bioinformatics 34, 3094–3100, doi:10.1093/bioinformatics/bty191 (2018).

90 Danecek, P. et al. Twelve years of SAMtools and BCFtools. GigaScience 10, giab008, doi:10.1093/gigascience/giab008 (2021).

91 Diesh, C. et al. JBrowse 2: a modular genome browser with views of synteny and structural variation. Genome Biology 24, 74, doi:10.1186/s13059-023-02914-z (2023).

92 Gel, B. & Serra, E. karyoploteR: an R/Bioconductor package to plot customizable genomes displaying arbitrary data. Bioinformatics 33, 3088–3090, doi:10.1093/bioinformatics/btx346 (2017).

93 Koren, S. et al. Canu: scalable and accurate long-read assembly via adaptive k-mer weighting and repeat separation. Genome Research 27, 722–736 (2017).

94 Cheng, H., Concepcion, G. T., Feng, X., Zhang, H. & Li, H. Haplotype-resolved *de novo* assembly using phased assembly graphs with hifiasm. Nat Methods 18, 170–175 (2021).

95 Manni, M., Berkeley, M. R., Seppey, M., Simão, F. A. & Zdobnov, E. M. BUSCO Update: Novel and Streamlined Workflows along with Broader and Deeper Phylogenetic Coverage for Scoring of Eukaryotic, Prokaryotic, and Viral Genomes. Molecular Biology and Evolution 38, 4647–4654, doi:10.1093/molbev/msab199 (2021).

96 Walker, B. J. et al. Pilon: An Integrated Tool for Comprehensive Microbial Variant Detection and Genome Assembly Improvement. PLOS ONE 9, e112963, doi:10.1371/journal.pone.0112963 (2014).

97 Flynn, J. M. et al. RepeatModeler2 for automated genomic discovery of transposable element families. 117, 9451–9457, doi:10.1073/pnas.1921046117 (2020).

98 Smit, A., Hubley, R. & Green, P. RepeatMasker Open-4.0. <http://www.repeatmasker.org> (2013-2015).

99 Bao, W., Kojima, K. K. & Kohany, O. Repbase Update, a database of repetitive elements in eukaryotic genomes. Mobile DNA 6, 11, doi:10.1186/s13100-015-0041-9 (2015).

100 Bruna, T., Hoff, K. J., Lomsadze, A., Stanke, M. & Borodovsky, M. BRAKER2: automatic eukaryotic genome annotation with GeneMark-EP+ and AUGUSTUS supported by a protein database. NAR Genomics and Bioinformatics 3, lqaa108 (2021).

101 Chen, S., Zhou, Y., Chen, Y. & Gu, J. fastp: an ultra-fast all-in-one FASTQ preprocessor. Bioinformatics 34, i884–i890, doi:10.1093/bioinformatics/bty560 (2018).

102 102 Wu, T. D., Reeder, J., Lawrence, M., Becker, G. & Brauer, M. J. in Statistical Genomics: Methods and Protocols (eds Ewy Mathé & Sean Davis) 283-334 (Springer New York, 2016).

103 Keller, O., Kollmar, M., Stanke, M. & Waack, S. A novel hybrid gene prediction method employing protein multiple sequence alignments. Bioinformatics 27, 757–763, doi:10.1093/bioinformatics/btr010 (2011).

104 Gabriel, L., Hoff, K. J., Brůna, T., Borodovsky, M. & Stanke, M. TSEBRA: transcript selector for BRAKER. 22, 566, doi:10.1186/s12859-021-04482-0 (2021).

105 Emms, D. M. & Kelly, S. OrthoFinder: phylogenetic orthology inference for comparative genomics. Genome Biology 20, 238, doi:10.1186/s13059-019-1832-y (2019).

106 Altschul, S. F., Gish, W., Miller, W., Myers, E. W. & Lipman, D. J. Basic local alignment search tool. Journal of Molecular Biology 215, 403–410, 10.1016/S0022-2836(05)80360-2 (1990).

107 Petit, E. et al. Linkage to the mating-type locus across the genus *Microbotryum*: insights into non-recombining chromosomes. Evolution 66, 3519–3533 (2012).

108 Hao, Z. et al. RIdeogram: drawing SVG graphics to visualize and map genome-wide data on the idiograms. PeerJ. Computer science 6, e251, doi:10.7717/peerj-cs.251 (2020).

109 Gu, Z., Gu, L., Eils, R., Schlesner, M. & Brors, B. circlize Implements and enhances circular visualization in R. Bioinformatics 30, 2811–2812, doi:10.1093/bioinformatics/btu393 (2014).

110 Ranwez, V., Douzery, E. J. P., Cambon, C., Chantret, N. & Delsuc, F. MACSE v2: Toolkit for the alignment of coding sequences accounting for frameshifts and stop codons. Molecular Biology and Evolution 35, 2582–2584, doi:10.1093/molbev/msy159 (2018).

111 Wertheim, J. O., Murrell, B., Smith, M. D., Kosakovsky Pond, S. L. & Scheffler, K. RELAX: Detecting Relaxed Selection in a Phylogenetic Framework. Molecular Biology and Evolution 32, 820–832, doi:10.1093/molbev/msu400 (2015).

112 Ranwez, V., Harispe, S., Delsuc, F. & Douzery, E. J. P. MACSE: Multiple alignment of coding sequences accounting for frameshifts and stop codons. PLOS ONE 6, e22594, doi:10.1371/journal.pone.0022594 (2011).

113 Yang, Z. PAML 4: Phylogenetic analysis by maximum likelihood. Molecular Biology and Evolution 24, 1586–1591, doi:10.1093/molbev/msm088 (2007).

114 Wickham, H. Ggplot2: Elegant Graphics for Data Analysis (2nd. ed.). (Springer Publishing Company, 2009).

115 Lindeløv, J. K. Mcp: An R package for regression with multiple change points. 10.31219/osf.io/fzqxv (2020).

116 Minh, B. Q. et al. IQ-TREE 2: New models and efficient methods for phylogenetic inference in the genomic era. Molecular Biology and Evolution 37, 1530–1534, doi:10.1093/molbev/msaa015 (2020).

117 Abascal, F., Zardoya, R. & Telford, M. J. TranslatorX: multiple alignment of nucleotide sequences guided by amino acid translations. Nucleic acids research 38, W7–13, doi:10.1093/nar/gkq291 (2010).

118 Edgar, R. C. MUSCLE: Multiple sequence alignment with high accuracy and high throughput. Nucleic acids research 32, 1792–1797, doi:10.1093/nar/gkh340 (2004).

119 To, T.-H., Jung, M., Lycett, S. & Gascuel, O. Fast dating using least-squares criteria and algorithms. Systematic biology 65, 82–97, doi:10.1093/sysbio/syv068 (2016).

120 Lopez-Villavicencio, M. et al. Multiple infections by the anther smut pathogen are frequent and involve related strains. PloS Pathogens 3, e176 (2007).

121 Vasimuddin, M., Misra, S., Li, H. & Aluru, S. Efficient architecture-aware acceleration of BWA-MEM for multicore systems. 2019 IEEE International Parallel and Distributed Processing Symposium (IPDPS), 314-324 (2019).

122 Manichaikul, A. et al. Robust relationship inference in genome-wide association studies. Bioinformatics 26, 2867–2873, doi:10.1093/bioinformatics/btq559 (2010).

123 Chang, C. C., et al. Second-generation PLINK: rising to the challenge of larger and richer datasets. Gigascience 4, 7, doi:10.1186/s13742-015-0047-8 (2015).

124 Untergasser, A. et al. Primer3--new capabilities and interfaces. Nucleic Acids Res. 40, e115 (2012).

125 Castle, A. J. & Day, A. W. Isolation and Identification of α -Tocopherol as an Inducer of the Parasitic Phase of Ustilago violacea Phytopathology 74, 1194–1200 (1984).

126 Gu, Z., Gu, L., Eils, R., Schlesner, M. & Brors, B. circlize implements and enhances circular visualization in R. Bioinformatics 30, 2811–2812, doi:10.1093/bioinformatics/btu393 (2014).

127 Kurtz, S. et al. Versatile and open software for comparing large genomes. Genome Biology 5, R12, doi:10.1186/gb-2004-5-2-r12 (2004).

